# A new mechanochemical vertex model with Ca^2+^ signalling, for apical constriction in neural tube closure

**DOI:** 10.1101/2024.12.31.630855

**Authors:** Abhishek Chakraborty, Timothy N. Phillips, Neophytos Christodoulou, Paris A. Skourides, Philip K. Maini, Ruth E. Baker, Katerina Kaouri

**Affiliations:** School of Mathematics, Cardiff University, Cardiff, CF24 4AG, UK; Department of Biological Sciences, University of Cyprus, Nicosia, CY-1678, Cyprus; Mathematical Institute, University of Oxford, Oxford, OX2 6GG, UK

## Abstract

Apical constriction during neural tube closure is driven by cell contractions which are preceded by asynchronous and cell-autonomous Ca^2+^ flashes, as demonstrated in recent experiments. Disruption of these Ca^2+^ signals and contractions leads to neural tube defects, such as anencephaly and spina bifida. A good understanding of the two-way mechanochemical coupling of Ca^2+^ signalling and mechanics remains elusive, while live-cell imaging is difficult. Thus, mathematical modelling is essential but existing models do not exhibit good agreement with experiments. We present two new mechanochemical vertex models of apical constriction during neural tube closure and simulate them using CelluLink, a new user-friendly open-source Python package for vertex modelling. The first, ‘one-way’ mechanochemical model only studies the effect of Ca^2+^ signalling on cell mechanics. It improves previous models, reproducing some key experimental observations, such as the reduction of the neural plate size to 2%-8% of its initial area. Other novel features of the one-way model is the incorporation of the surface ectoderm and of the experimental amplitude and frequency profiles of the Ca^2+^ flashes. Furthermore, guided by experiments, the damping coefficient of the vertices and cell-cell adhesion are modelled as functions of the actomyosin concentration and cell size. Furthermore, we develop a ‘two-way’ model which improves the one-way model by capturing the two-way coupling between Ca^2+^ signalling and cell mechanics, through the incorporation of stretch-sensitive Ca^2+^ channels. These channels enable cells to sense mechanical stimuli and encode them into Ca^2+^ signals. In the two-way model, the Ca^2+^ flash frequency and amplitude profiles are model outputs and are not inputs as in the one-way model. Finally, we use both models to propose a series of hypotheses for future experiments.

**Author summary:** As a baby is growing in the womb, its neural tube closes to form the brain and the spinal cord. During neural tube closure, cells are contracting in a ratchet-like way while experiencing a ‘choreography’ of Ca^2+^ flashes. If the Ca^2+^ flashes or the contractions go wrong, serious birth defects like spina bifida and anencephaly may arise. Understanding how Ca^2+^ flashes and contractions work together is complex, especially since studying living cells is challenging. To address this challenge, we developed two new mathematical models. The first model captures how Ca^2+^ flashes affect contractions, improving previous models and accurately capturing some experimental results. For example, it incorporates recent experimental measurements of the amplitude of Ca^2+^ flashes’ (brightness) and their frequency (how frequently the flashes appear). The second model builds on the first model by additionally capturing the effect contractions have on the Ca^2+^ flashes. We capture this two-way coupling by enabling cells to sense mechanical stimuli through stretch-sensitive Ca^2+^ channels. In this case, the amplitude and frequency of the Ca^2+^ flashes arise as outputs. Both models inform future experiments that will further elucidate embryo malformations.

## Introduction

A critical event during vertebrate embryogenesis is the formation of the central nervous system, with the initial crucial step being the formation of the neural tube. The neural tube originates from the dorsal ectoderm, a layer of cells (Fig 1a), which is the precursor to the brain and spinal cord. Defects in neural tube formation, like spina bifida and anencephaly, are a leading cause of birth defects, occurring in 0.5-2 per 1000 births [1]. Neural tube defects can lead to neurological impairments, such as paralysis and difficulties with bladder and bowel control in cases of spina bifida, or fatality in cases of anencephaly. Therefore, a thorough understanding of the morphogenetic events that drive neural tube formation is essential for comprehending and preventing these defects.

Given the significance of neural tube formation in embryonic development, model systems have been extensively used to investigate the cellular, molecular, and biophysical events involved [2–5]. Neural tube formation progresses through distinct stages. Initially, the dorsal ectoderm differentiates into the neural plate, followed by tissue thickening through cell elongation. Subsequently, the neural folds at the lateral edges elevate and bend towards the midline [6]. This medial movement is driven by tissue folding, mediolateral narrowing, and anteroposterior elongation of the neural plate. Finally, the neural folds converge and fuse at the midline, completing neural tube formation through cell protrusion-mediated fusion of the non-neural ectoderm [7, 8].

The transformation of the flat neuroepithelium into the neural tube is driven by intrinsic forces generated by morphogenetic events within the neural plate and extrinsic forces affected by the behaviour of neighbouring tissue [9, 10]. One key morphogenetic process necessary for neural tube closure (NTC) is apical constriction (AC). AC is a process that converts columnar cells into wedge-shaped cells by reducing their apical surface area. In Fig 1a we show the posterior NP and the Anterior NP during AC; the ectoderm is also shown. AC generates contractile forces that drive epithelial folding and is critical for bending the neuroepithelium during neurulation [11–13].

AC during NTC is regulated by cell-autonomous and asynchronous Ca^2+^ flashes [11]. A Ca^2+^ flash (also referred to as a Ca^2+^ transient, Ca^2+^ pulse or Ca^2+^ oscillation) occurs when the cytosolic Ca^2+^ concentration oscillates between a low (inactivated) to a high Ca^2+^ concentration (activated) state [11, 14, 15] (see Fig 1c). Cell-autonomous Ca^2+^ flashes (see Fig 1b, Fig 1c) regulate medio-apical actomyosin contraction and monotonic apical tissue surface reduction [11] (see Fig 1c). The apical surface of the anterior NP contracts to within 2% to 8% of its initial area [9], while the Ca^2+^ flash amplitude and frequency increase by more than 50% of their initial values. The AC process in NTC takes approximately 40 to 60 minutes in *Xenopus* [9, 11]. Note that no model of NTC [14, 16] has reproduced this key behaviour to date; we set out to reproduce this here, along with several other experimental findings.

**Fig 1.**
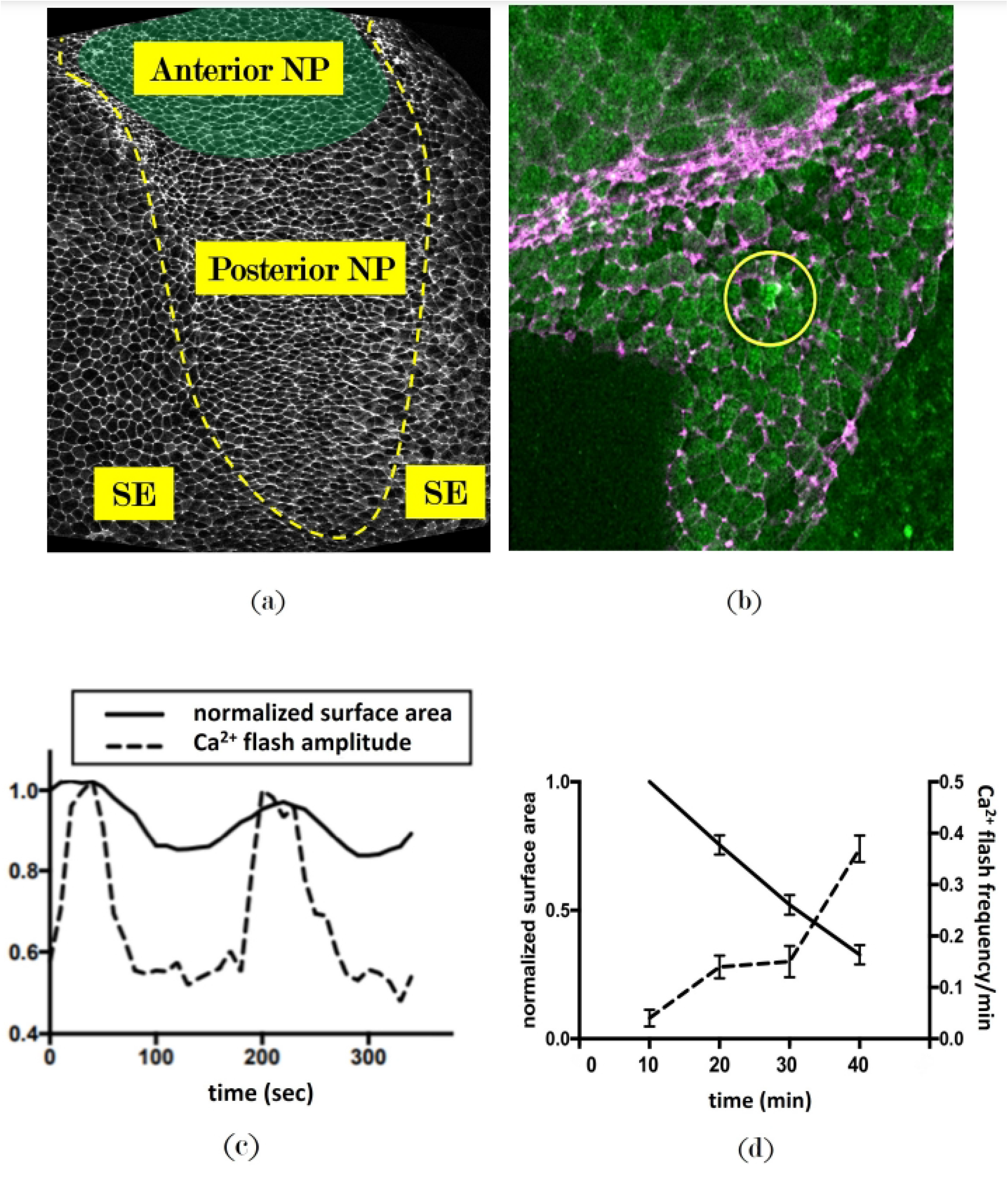
Ca^2+^ flashes in an embryo undergoing NTC. (a) Extract from a *Xenopus* embryo at the onset of NTC. The dashed yellow line marks the border between the surface ectoderm (SE) and neural plate (NP); the anterior NP and the posterior NP are shown. Cell boundaries were tracked using Phalloidin-488; (b) During NTC, anterior NP cells display Ca^2+^ flashes (bright green). Cell cortices (purple) were tracked using mem-GFP and cytosolic Ca^2+^ levels (green) were tracked using GECO-RED. Images (a) and (b) were taken using a ZEISS LSM710 confocal microscope; (c) Normalised Ca^2+^ flash amplitude and normalised apical surface area of a single cell during AC; (d) Normalised Ca^2+^ flash frequency and normalised apical surface area, averaged over 10 cells, during AC. Image source: [11, 17].

Ca^*2+*^ flashes *precede* contractions in single epithelial cells (see Fig 1c) [11, 14, 17], i.e., there is a latency between the occurrence of a Ca^2+^ flash and the onset of cell contraction. The contractions, characteristic in AC [18], influence the behaviour of neighbouring cells, strongly suggesting that mechanical coupling is essential for translating cell-autonomous events into global tissue shape changes [19]. Despite the clear role of mechanotransduction in AC, understanding how mechanical coupling directly affects cell behaviour during this process remains challenging with current experimental methods.

Therefore, mathematical modelling and simulations are essential. There are, however, very few models of Ca^2+^ signalling in AC [14, 16, 17], and they only capture a small number of experimental behaviours. The continuum models developed by Kaouri et al [16, 17] build on early continuum models that couple Ca^2+^ dynamics to cell mechanics and replace the hypothetical bistable Ca^2+^ release [20–23] with the experimentally validated IP_3_-mediated Ca^2+^ dynamics of the Atri model [24]. In this model, a advection-diffusion-reaction system for Ca^2+^ concentration is coupled to a force balance equation for the tissue, which is assumed to be a linear, viscoelastic material. The ectoderm is also modelled in [16] and the Ca^2+^ flashes are implemented as a random distribution of Ca^2+^ flashes. A novel contribution in [17] is deriving a ‘stretch activation’ Ca^2+^ flux as a bottom-up contribution from stretch-sensitive Ca^2+^ channels (SSCCs) on the cell membrane [25–28] ; we are also going to incorporate SSCCs in our modelling here. The model in [16] reproduces some important experimental features, such as the monotonic reduction of the apical surface area of the anterior neural plate. Both models demonstrate that mechanical effects can cause Ca^2+^ oscillations to vanish, which implies information loss and corresponds to failure of AC and other key embryogenic processes. Continuum models, however, are unable to resolve individual cells and capture crucial cell-level behaviours, such as the ‘pulsed’ cell contractions or cell shape changes which are crucial in AC [18].

Hence, cell-based models are needed; we choose to work with vertex models since cells can be modelled as polygons, which closely resemble real cell shapes (see Fig 1a, b). Vertex models have been developed for a wide range of applications, ranging from the study of *Drosophila* wing disc formation [29] to the understanding of tumour growth [30] and wound healing [31]. The movement of cell vertices is governed by mechanical forces, allowing the models to capture detailed cell-level behaviours such as changes in cell shape, division, and migration.

Suzuki et al [14] developed the first and only mechanochemical vertex model that studies the impact of Ca^2+^ flashes in AC during NTC, in the *Xenopus* neural plate. The model is in agreement with some experimental observations [14]. For instance, their model reproduces the relaxation events following the Ca^2+^-induced contractions (Fig 1b) and that Ca^2+^ flashes accelerate AC. Furthermore, their model suggests that cell-autonomous Ca^2+^ flashes are able to reduce tissue size more effectively than multicellular Ca^2+^ signals. For simplicity, we will refer to their model as the Suzuki model for the remainder of this work.

The Suzuki model has, however, some limitations. First, crucially, the model does not consider the surface ectoderm. During AC, the contraction of the anterior NP exerts a force on the layer of cells of the surface ectoderm, deforming and displacing them. Consequently, the ectoderm exerts a resistive force on the anterior NP, significantly influencing morphogenesis during NTC [9] and needs to be included in the modelling. Moreover, the Suzuki model does not incorporate the increasing frequency and amplitude of Ca^2+^ flashes [17]. Also, it is assumed that a Ca^2+^ flash triggers cell contraction immediately instead of the Ca^2+^ transient *preceding* contraction. Finally, and very importantly, the model does not account for the two-way feedback between Ca^2+^ and cell mechanics. As a result, the model is not able to reproduce the 2% to 8% reduction in the initial area of the anterior NP over the 40-60 minutes that AC lasts; the simulated reduction is only around 20 − 30%.

In this work, we use the Suzuki model as a foundation for building an improved mechanochemical vertex model. We develop a new mechanochemical model of AC during NTC that captures the two-way coupling between Ca^2+^ and cellular mechanics which agrees *quantitatively* with the experimentally observed contraction of the apical surface area and with several other experimental findings. We will refer to this model as the two-way mechanochemical model or **the two-way model**.

We first construct a one-way mechanochemical model (or **the one-way model**) as an important intermediate step towards building the two-way model. We use the one-way model to study only the effect of Ca^2+^ on cell mechanics and tissue behaviour. Then, in the two-way model, the effect of Ca^2+^ on cell mechanics is modelled in the same manner as the one-way model but to capture the effect of cell mechanics on Ca^2+^ signalling, we additionally incorporate SSCCs [22] which sense mechanical deformations of a cell, leading to Ca^2+^ flashes. Although the two-way model is more complete than the one-way model, the latter is very useful for studying the effect of Ca^2+^ on cell mechanics in the absence of the two-way coupling. For example, incorporating more sophisticated Ca^2+^ dynamics, such as IP_3_-mediated Ca^2+^ dynamics, into the two-way model could begin by first integrating these dynamics into the simpler one-way model; the insights gained could then be used to easily inform modifications to the two-way model.

### Vertex models

Since both the one-way and two-way models are vertex models, we briefly outline the basics of vertex models in this section. In vertex models, a tissue is represented by a collection of non-overlapping connected polygons whose vertices are free to move and each polygon corresponds to a cell, i.e., the tissue is represented as a convex polygonal partitioning of the plane (Fig 2).

Following the standard formulation of vertex models [14, 29, 32–34], the total potential energy of the tissue, *U*, is defined as follows:

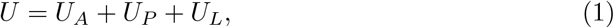

where *U*_*A*_ is the potential energy of the cell areas, *U*_*P*_ is the potential energy of the cell perimeters and *U*_*L*_ is the potential energy of the cell-cell junctions. Also, as is typical of vertex models [14, 31–33], we define the force on the *i*^*th*^ vertex in the tissue, **F**_**i**_, as follows:

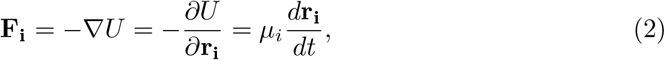

**Fig 2.**
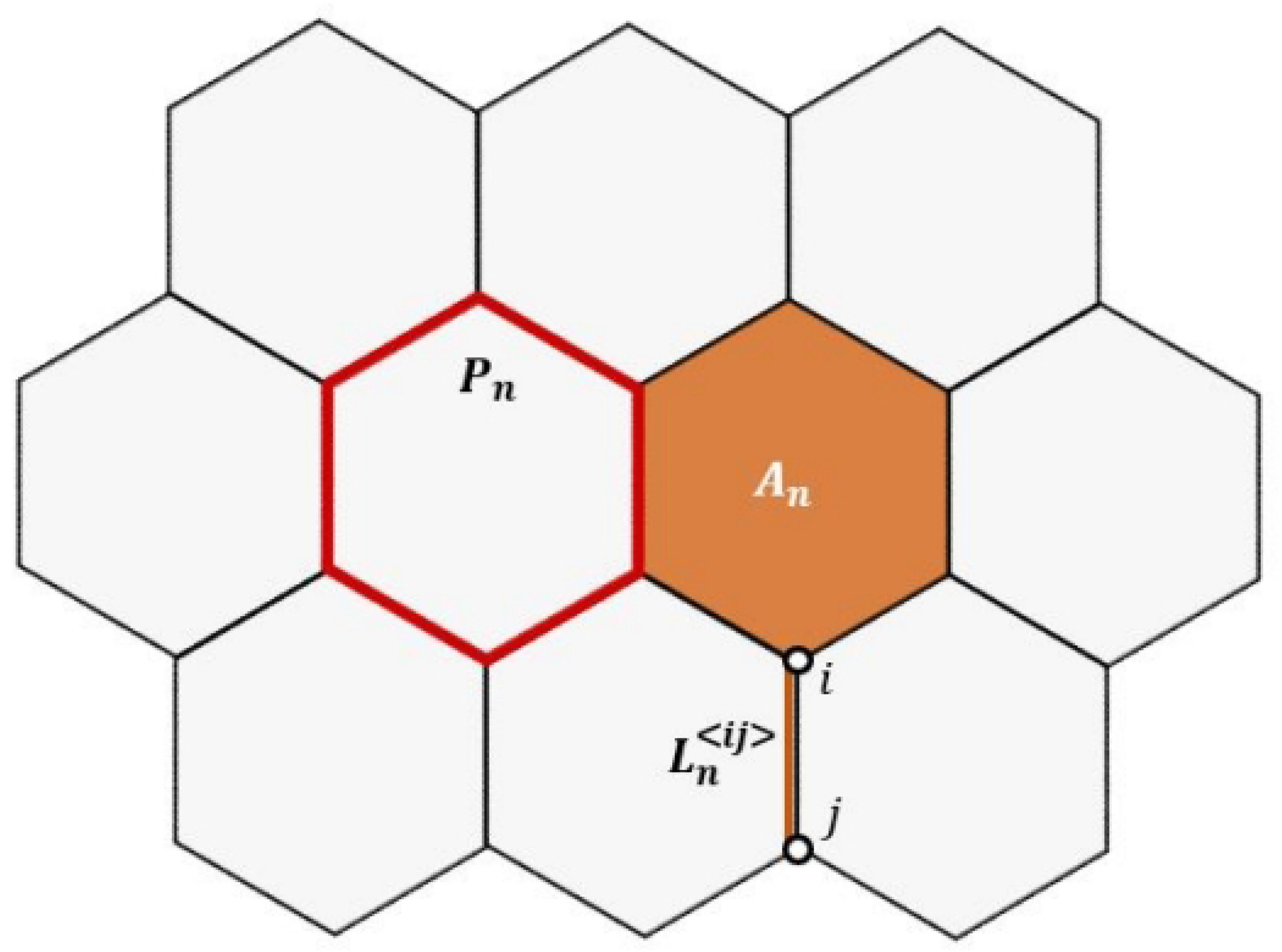
Schematic representation of cells in the vertex model. The three physical attributes, cell area (*A*_*n*_), cell perimeter (*P*_*n*_), and edge length 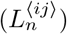, determine the mechanical potential energies: *U*_*A*_, *U*_*P*_, and *U*_*L*_, respectively. The edge 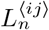 connects vertices *i* and *j*.

where **r**_**i**_ = (*x*_*i*_, *y*_*i*_) is the position of the *i*^*th*^ vertex, and *µ*_*i*_ is the damping function of the *i*^*th*^ vertex, emcompassing the effect of all dissipative effects –viscosity and friction between neighbouring cells as well as between the cells and the substrate and the extracellular matrix. This *overdamped limit* is still a good assumption for biological systems, since the inertial effects (acceleration) are typically several orders of magnitude smaller than the effects arising from cell-cell interactions [31], as cells move in dissipative environments with very small Reynolds number [38]. Vertex models are capable of capturing many key features of real epithelial tissues [14, 35–37].

While the exact definitions of *U*_*A*_ and *U*_*P*_ vary across studies, they typically increase quadratically as the cells deviate from their natural area and perimeter, respectively. Specifically, the following expressions are usually used:

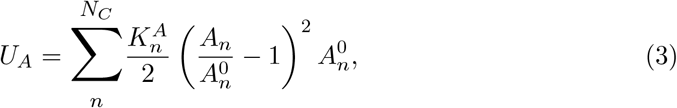

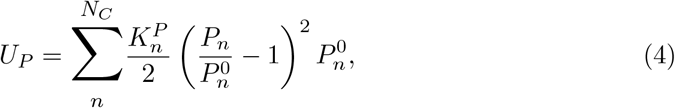

where 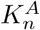 is the area elasticity coefficient, *A*_*n*_ is the area of the cell, 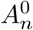 is the natural area of the cell, 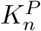 is the perimeter elasticity coefficient, *P*_*n*_ is the perimeter of the cell, 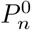 is the natural perimeter of the cell, and *N*_*C*_ is the number of cells in the tissue. The physical attributes of a cell, area (*A*_*n*_), perimeter (*P*_*n*_), and edge length 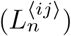, are shown in Fig 2. 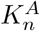 and 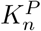 are a measure of the concentration of actin in the medioapical cortex and at the cell-cell junctions, respectively.

Vertex models minimise the total energy of the tissue, as dictated by the laws of thermodynamics, considering contributions from factors like tension and adhesion at the cell-cell junctions and area and perimeter elasticities [14, 29, 32–34].

### Mechanochemical vertex models and the Suzuki model

Here, we give a brief summary of the Suzuki vertex model since it is the foundation of our models. The Suzuki model inherits from the general vertex modelling framework the governing equations (1)-(4); we will also use these equations in the one-way model and in the two-way model.

In the Suzuki model, *U*_*L*_ captures the effect of line tension along a cell edge and is given by the standard expression formula

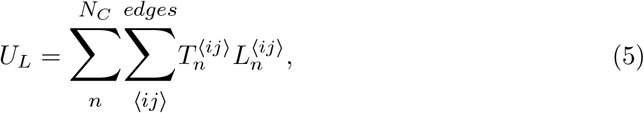

where *T*_*n*_, the line tension along a cell edge, is expressed as a sum of baseline tension and a Ca^2+^-dependent tension, *ξ*:

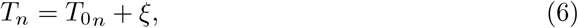

where 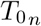 is the baseline value of line tension and characterizes the contractile action of myosin motors attached to the junctional actin network.

Note that in the new mechanochemical models, we will incorporate an adhesion force in addition to line tension to provide a more accurate description of the biology, specifically the role of adherens junctions [37, 39, 40], which results in a different expression for *U*_*L*_–please see the following section.

A novel feature of the Suzuki model is the modelling of the ratchet-like behaviour of the cells during AC. During AC, actin filaments accumulate in the apical cortex, providing a substrate for the binding of myosin molecules [18]. The motor activity of the myosin molecules causes actin filament contraction, which results in a reduction of the cell’s apical surface area. The relaxation of the apical surface area results from the exhaustion of the myosin motors, which detach from the actin filaments, allowing for the relaxation of the actin network back to a ‘rest area’ [18]. If the actin network is reassembled during contraction, the network can no longer relax to its previous rest area and instead settles into a smaller rest area. The repeated assembly and breakdown of the actomyosin bundles result in pulsed contractions of the apical area. Thus, the *ratchet-like mechanism* of contraction and stabilization arises [18].

The Suzuki model represents the ratchet-like mechanism as two first-order, linear ordinary differential equations that track a cell’s natural area, 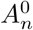, and natural perimeter, 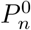, respectively:

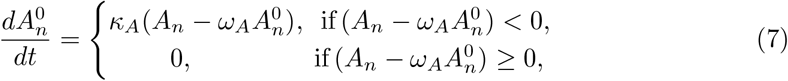

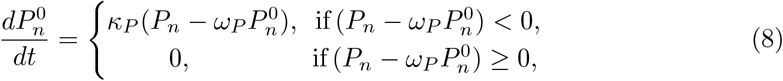

where *κ*_*A*_ and *κ*_*P*_ are rate constants. According to these equations, 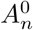 and 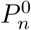 are reduced when the cell’s area, *A*_*n*_, and perimeter, *P*_*n*_, drop below predefined thresholds, *ω*_*A*_ and *ω*_*P*_, respectively.

### The one-way mechanochemical model

We now present the one-way model, a new mechanochemical vertex model for the constriction of the anterior NP during the AC phase of NTC in *Xenopus*. The model improves the Suzuki model in several ways, achieving quantitative agreement to many experimental findings. Like the Suzuki model, the one-way model captures the effect of Ca^2+^ on cellular mechanics but not vice versa.

We begin by describing the computational domain over which the governing equations for the one-way and for the two-way model are solved. This is followed by the governing equations and modelling assumptions. The assumptions are motivated, leveraging all the experimental data available to date. Finally, we simulate the one-way model and discuss its behaviour.

### Computational domain

We model the anterior NP as hexagon-shaped (see Fig 3), which reasonably approximates the anterior NP (see Fig 1a). We assume that the anterior NP comprises 271 cells, which closely matches the number of cells studied experimentally in [9, 11].

**Fig 3.**
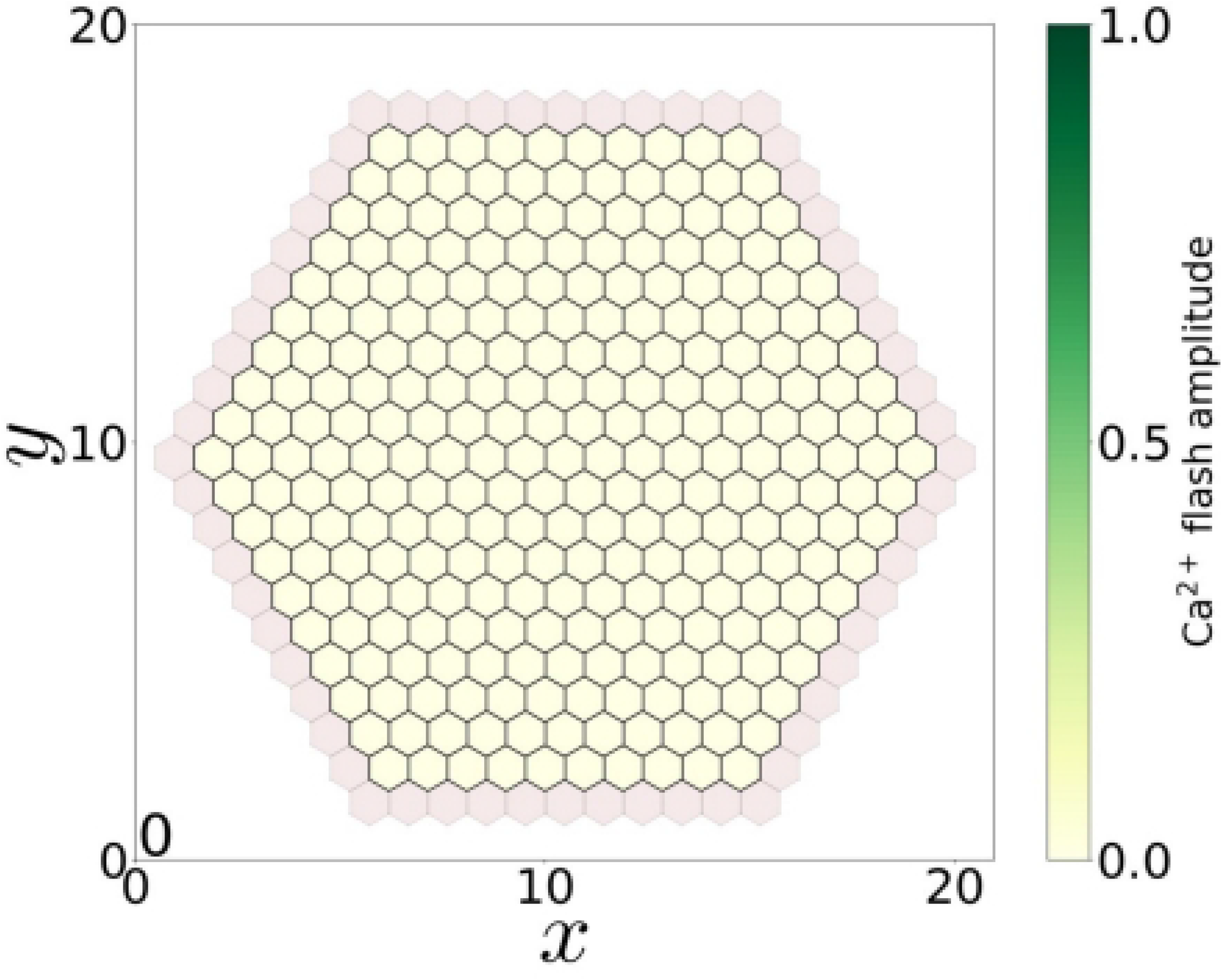
The initial configuration of the simulated anterior NP and virtual cells in the one-way model. The apical surface of the anterior NP cells (solid yellow) and the virtual cells representing the surface ectoderm layer (pink), at the start of the simulation.

The initial configuration of the anterior NP and surface ectoderm is a hexagonal honeycomb lattice, with cells arranged in a hexagon-shaped tissue. Initially, each cell is assumed to be a regular hexagon with area 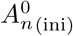.

The polygonal shape of the epithelial plate cells (Fig 1a) can be modelled accurately using a two-dimensional (2D) vertex model. Although the apical surface of the anterior NP bends out of the 2D plane during NTC, a 2D model is still sufficient to explore the two-way coupling between Ca^2+^ signalling and cellular mechanics [26–28, 42]. Note that the Suzuki model [14] is also two-dimensional.

A crucial component we incorporate in the model is the surface ectoderm. Although the surface ectoderm comprises multiple cell layers, we model it as a single layer of 60 ‘virtual’ cells (Fig 3). This is sufficient to simulate the ectoderm’s mechanical effects on the NP’s deformation. During the simulation, the outer edge of the surface ectoderm layer, which is stiffer [43], remains fixed in place, while the inner edge is pulled inwards by the contraction of the NP. The deforming virtual cells exert an elastic force on the anterior NP which increases as the cells deform further. While the contraction of the anterior NP cells is driven by Ca^2+^ flashes, the cells of the surface ectoderm do not experience any Ca^2+^ flashes [9, 11].

Typically, vertex models, including the Suzuki model, implement topology changing events like T1 and T2 processes [36, 44, 45] to resolve small cell edges and prevent instabilities. These events correspond to cell-cell intercalation (neighbour exchanges) and cell death. Since the anterior NP does not experience these events during AC [9], T1 and T2 processes are not implemented in our model.

### Governing equations and modelling assumptions

The one-way model, like the Suzuki model [14], inherits, from the general vertex modelling framework, Equations (1)-(4) which describe the potential energies for cell areas, perimeters, and vertex dynamics. Equations (5) and (6) introduced by the Suzuki model to model the ratchet-like mechanism will be modified accordingly. Furthermore, we introduce several novel features, equations, formulae and underlying assumptions which are also presented and justified in detail below. Details of the parameter estimation can be found in Appendix S1A. As is typical for vertex models, all values are in non-dimensional or arbitrary units (a.u.).

### Ca^2+^ flash frequency and amplitude

The frequency and amplitude profiles for the Ca^2+^ flashes are fitted to the experimental data in Christodoulou and Skourides [11], as follows:

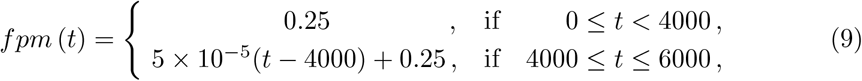

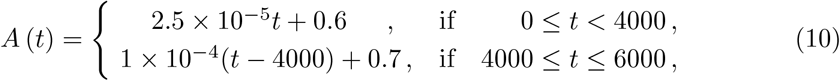

where *fpm* (*t*), the number of flashes per minute per cell, determines the frequency of Ca^2+^ flashes and *A* (*t*), the flash amplitude, represents the cytosolic Ca^2+^ concentration. At each time step during the simulation, the values of *fpm* and *A* are updated according to Equations (9) and (10) to accurately capture the frequency and amplitude profiles of the Ca^2+^ flashes.

### Probability of cell activation

#### Assumption 1.1.

*When a cell experiences a Ca*^*2+*^ *flash, its Ca*^*2+*^ *level is elevated and it becomes activated for a time period. The activation period is followed by a refractory interval when its Ca*^*2+*^ *level returns to baseline. After the refractory period lapses, the cell has a non-zero probability of experiencing a Ca*^*2+*^ *flash and becoming activated again*.

In the Suzuki model, it is assumed that a cell can experience sustained elevation in Ca^2+^ levels without an intervening refractory period. However, prolonged elevation of Ca^2+^ levels can lead to the failure of NTC [11], highlighting the need to model the refractory interval.

Based on experimental data [11, 17], we estimate the activation time interval (*τ*) and the refractory interval to be twice the inactivation interval of a cell (2*τ*). The activation and inactivation intervals form the period of the Ca^2+^ oscillation, *T*_*osc*_ = 3*τ* .

The activation probability of a cell, *p*_*c*_, is updated at the start of every *T*_*osc*_-sized interval. If the cell does not experience a Ca^2+^ flash, we assume that the cell is maintained in an inactive state for time interval *T*_*osc*_, after which there is a non-zero probability that the cell will experience a Ca^2+^ flash and become activated again.

Let *T*_60_ denote the non-dimensional time interval corresponding to 60 minutes, the duration of the AC process. If we consider *T*_60_ to be divided into smaller intervals of duration *T*_*osc*_, then the probability of cell activation, *p*_*c*_, is defined as:

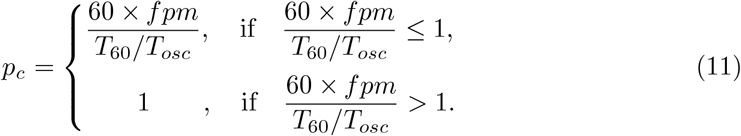

### Ca^2+^-induced elevation in line tension

#### Assumption 1.2.

*Upon cell activation, the line tension of each cell edge increases sigmoidally over the course of the activation*.

In the Suzuki model, it is assumed that a Ca^2+^ transient triggers cell contraction instantaneously and that *ξ* is a constant. However, Ca^2+^ flashes precede cellular contractions [11, 14, 17], i.e., there is a latency between the two events. This latency stems from the time required for the entire sequence of events, from the binding of Ca^2+^ to Ca^2+^-binding proteins to the subsequent contraction of the actomyosin complexes [46]; this suggests that *ξ* should be modelled as a sigmoidal response to the Ca^2+^ flash amplitude.

To determine the shape of the sigmoidal function, we assume that a larger Ca^2+^ flash amplitude causes a steeper increase in line tension, as the amplitude of a Ca^2+^ flash drives myosin activity and cell contraction [11]. During AC, actomyosin accumulates in the apical cortex of NP cells [11, 14], requiring progressively higher Ca^2+^ levels for activation. If an insufficient amount of actomyosin is activated, the resulting increase in line tension cannot overcome the elastic forces of the accumulated actomyosin. This necessitates greater Ca^2+^ flash amplitudes to initiate sufficient contraction of the cell’s apical surface.

Summarising the above, we represent *ξ* as follows:

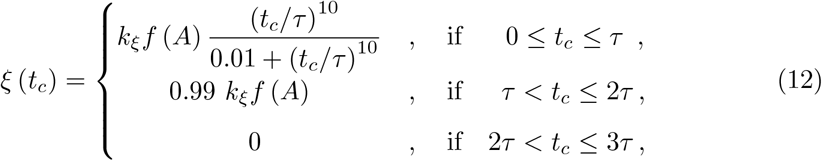

and

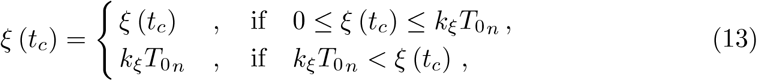

where *A* is the Ca^2+^ flash amplitude, *τ* is the activation interval of the cell, *t*_*c*_ represents the time elapsed since the activation of the cell, *k*_*ξ*_ is the scaling factor for 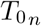, and *f* (*A*) represents the actomyosin activation as a function of Ca^2+^ flash amplitude, as follows:

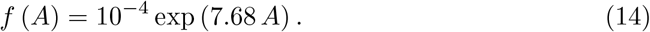

The sigmoidal function in Equation (12) is a Hill function, commonly employed to study the kinetics of many enzyme-catalyzed reactions and transporter-mediated processes [47]. Due to lack of experimental data for the Hill coefficient, we performed a parameter sweep and found that when the Hill coefficient is equal to 10 we obtain the closest match to the experimental cell contraction profile in [11, 17].

The cell is activated by a Ca^2+^ flash at *t*_*c*_ = 0 and the activation lasts for a duration *t*_*c*_ = *τ* . The Hill function reaches 99% at *t*_*c*_ = *τ*, i.e, when the cell transitions from a high to the baseline Ca^2+^ state. After the cell returns to the baseline Ca^2+^ state, the myosin motors remain active for a time, and we assume that *ξ* stays constant for a time *τ* before eventually dropping to 0, due to the exhaustion of myosin. Following this, the value of *ξ* is assumed to be 0, again, for a time interval *τ* . We estimate these time intervals from the experimental data in [11, 17]. In Equation (13), if *ξ* exceeds 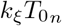 we set 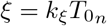 because actomyosin at the cell-cell junctions cannot contract indefinitely [48].

The precise mechanism governing the relationship between Ca^2+^ flash amplitude and actomyosin activation is unknown. However, there is a correlation between NTC velocity and Ca^2+^ flash amplitude [11], indicating that higher Ca^2+^ flash amplitudes may enhance actomyosin contractility. So, we use an exponential function for *f* (*A*) that fits the known data points 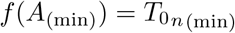 and 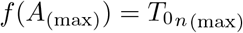. Since 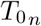 increases with the accumulation of actomyosin during AC, this approach uses 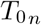 as a measure of the actomyosin level.

As AC commences, the actomyosin level in the apical cortex is at its lowest [9]; we take 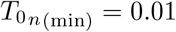. At this stage, a Ca^2+^ flash with an amplitude of *A*_(min)_ = 0.6 is sufficient to activate the existing actomyosin. Towards the end of AC, the actomyosin level in the apical cortex reaches its peak and we take 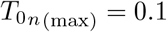. At this stage, a Ca^2+^ flash amplitude of *A* = 0.6 can activate only a fraction of the total actomyosin—to fully activate the increased amount of actomyosin accumulated in the apical cortex, a larger Ca^2+^ flash amplitude is required; we, thus, take *A*_(max)_ = 0.9.

### Adhesion force

#### Assumption 1.3.

*When the length of a cell edge decreases below a certain threshold, an adhesion force that opposes line tension and increases edge length is generated. This force is negligible for large edge lengths but dominates line tension for small edge lengths*.

As the apical surface area of a cell decreases, compression of the cytosol raises internal pressure, exerting a force that tends to increase cell edge length [37, 39]. Additionally, adherens junctions may also generate an adhesion force that counteracts the line tension [40]. Apical adherens junctions play a pivotal role in mediating mechanical coupling between epithelial cells. These junctions contain adhesion receptors (e.g., E-Cadherin) that facilitate homophilic adhesion between the cells of the epithelial layer [40]. The application of tension triggers the formation of adhesion complexes, increasing the size of adherens junctions and strengthening cell-cell adhesion [49–51].

Furthermore, elevated Ca^2+^ activity can enhance the function of adhesion receptors like E-Cadherin, further contributing to the elevation of adhesion strength within adherens junctions [52]. Therefore, for small edge lengths, an adhesion force that opposes further area reduction originates from the cell’s internal pressure, adherens junctions, or both, opposing tension and increasing interfacial contact between cells. To preserve cell integrity during morphogenesis, the adhesion force should be negligible compared to the line tension for larger edge lengths but has to dominate the line tension as the edge length decreases. We, thus, model the adhesion force along a cell edge, 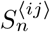, as follows:

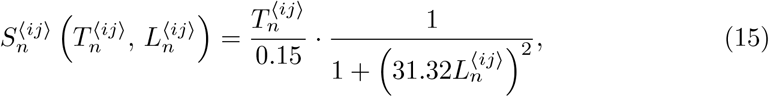

where 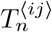 is the line tension along the cell edge, *n* denotes the cell index, and *ij* denotes the edge connecting vertices *i* and *j*.

We subtract 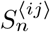 from 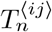 and integrate the result with respect to 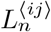, which yields the expression for the potential energy of a cell edge. Therefore, the total potential energy of the cell edges, *U*_*L*_, is defined as:

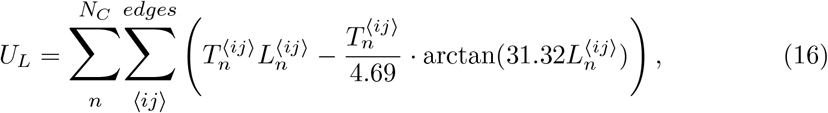

where *edges* and *N*_*C*_ indicate summation over all the edges of a cell and over all the cells in the tissue, respectively. The full derivation for 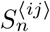 and *U*_*L*_ can be found in Appendix S1B.

### Damping function

#### Assumption 1.4.

*As the apical area of a cell decreases, the damping experienced by its vertices increases*.

In vertex models, the damping coefficient of the vertices, *µ*_*i*_, are typically taken to be constant (and equal) [31–33]; similarly in the Suzuki model [14]–see (2). However, experimental evidence indicates that vertex motion becomes increasingly damped as AC progresses [11, 14, 18]. While cells are contracting, actomyosin complexes accumulate in the apical cortex of the anterior NP cells due to the effect of the Ca^2+^ flashes [11, 14]. This accumulation of cortical actomyosin inhibits vertex movement [18], which we model as an increase in *µ*_*i*_. This is achieved by assuming that the damping effect of actomyosin varies inversely with 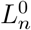, the natural length of the cell, so that *µ*_*i*_ is given by

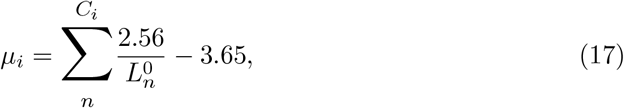

where *C*_*i*_ is the set of cells sharing the *i*^*th*^ vertex and

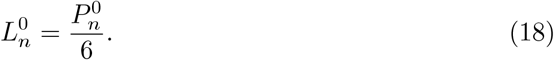

The constants in Equation (17) arise from linearly interpolating data at the start of the simulation and when a cell has contracted to its minimum size. The full derivation can be found in Appendix S1C.

### Ratchet-like mechanism

#### Assumption 1.5.

*If the apical area of a cell decreases below a predefined threshold, it cannot recover to its previous value when the forces are relaxed*.

Since the rate of reduction of 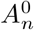 and 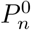 in the Suzuki model (Equations (7)-(8)) is directly proportional to 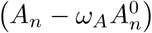 and 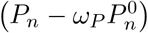, respectively, their ratchet-like mechanism does not activate for small cell sizes. Therefore, we modified the equation for updating 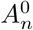 as follows:

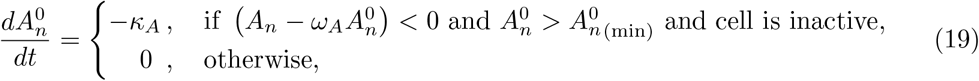

where *κ*_*A*_ is the constant contraction rate of 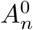 and 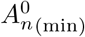 is the minimum value of 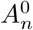. In the absence of relevant quantitative data, we assume a constant reduction rate, for simplicity. We no longer need an ODE for 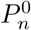; since cells are modelled as regular hexagons, we determine 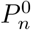 using the relationship between the area and perimeter of a regular hexagon:

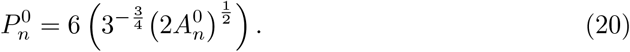

### Growth laws for the elastic constants and line tension

#### Assumption 1.6.

*Mechanical parameters grow linearly if a cell is in its refractory interval and has non-zero Ca*^*2+*^*-induced elevation in line tension, ξ >* 0.

Suzuki et al [14] modeled 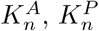, and 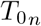 as constants, as it is the usual practice in vertex models. In the one-way model we will model 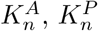, and 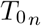 to be increasing with time in order to capture the increase of contractility and tension in NP cells due to actomyosin accumulation and rearrangement in the cell cortex, following activation by a Ca^2+^ flashes, as AC progresses [11, 14, 18, 53, 54].

The pharmacological elevation of cytosolic Ca^2+^ levels leads to synchronous and continuous cellular contractions, which interrupts AC and results in the failure of NTC [11]. This suggests that a refractory interval, characterized by low Ca^2+^ levels, is essential for activating the ratchet-like mechanism, which corresponds to increasing 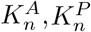, and 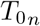. Based on the following experimental evidence:

i. Actomyosin bundles can be rearranged by the application of a mechanical force [53, 54],
ii. Ca^2+^ flashes occur just before a cell contraction pulse but never during the stabilization step [11],

we assume that the Ca^2+^-induced elevation in line tension causes the rearrangement of actomyosin bundles, and this rearrangement - indicated by permanent changes in 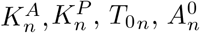, and 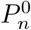-can only occur when the cell is not activated.We, thus, assume growth laws for 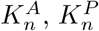, and 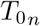, as follows:

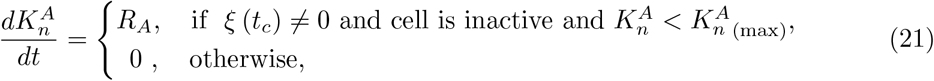

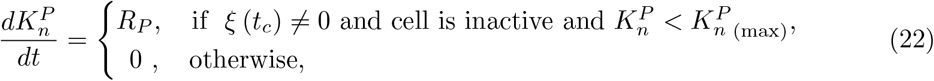

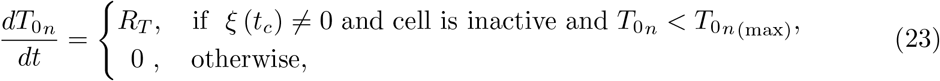

where *R*_*A*_, *R*_*P*_, and *R*_*T*_ are the constant growth rates of 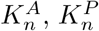, and 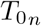, respectively, and 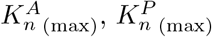, and 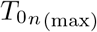 are the maximum values of of 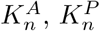, and 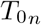, respectively. In the absence of quantitative data, we assume constant growth rates.

### Parameters and units

Since there is no straightforward method that relates the parameters of the vertex model (in non-dimensional units) to their experimentally measurable counterparts (in dimensional units), we estimate the parameter values using insights from simulation results, and experimental data from [9, 11, 14, 17, 43].

The rationale for the choice of parameter values in Table 1 is provided in Appendix S1A. As is typical for vertex models, all values are in non-dimensional or arbitrary units (a.u.). However, since successful apical constriction requires demonstrating sufficient area reduction within a specific timeframe, we establish the following relationship between non-dimensional and dimensional time units.

Since the apical surface of the anterior NP must contract to between 2% and 8% of its initial areas over the course of AC, which lasts between 40 and 60 minutes [9], we equate *t* = 6000 a.u. to 60 minutes. Therefore, *t* = 100 in the simulations corresponds to 1 minute. Because we are only concerned with the percentage reduction in area, all figures in this study use non-dimensional length units.

Similar to prior experimental studies [11, 17] on the mechanochemical coupling between Ca^2+^ and mechanics, we represent Ca^2+^ flash amplitude using non-dimensional units. In those studies, the Ca^2+^ flash amplitude was normalized by dividing all values by the maximum amplitude observed during the AC phase of NTC. Like those studies, our focus is on the relative strength of the Ca^2+^ flash amplitude rather than the exact dimensional values.

### Simulations

The one-way model consists of Equations (1)-(4), Equation (6) and Equations (9)-(23). We simulate the model using CelluLink, over the domain described above, for the parameter values given in Table 1. The ODEs were solved using the forward Euler method with time step *δt* = 0.2. We have developed CelluLink as a new open-source, user-friendly Python package for vertex modelling [41]. CelluLink can generate an hexagonal honeycomb lattice of cells arranged in both hexagon-shaped and rectangle-shaped tissues and can be used to study many other cell-based challenges.

Fig 4(a),(b),(c),(d) visualise the reduction of the apical surface area of the anterior NP, as simulated by the one-way model, for *t* = 0, 12, 24, 60 minutes, respectively.

**Fig 4.**
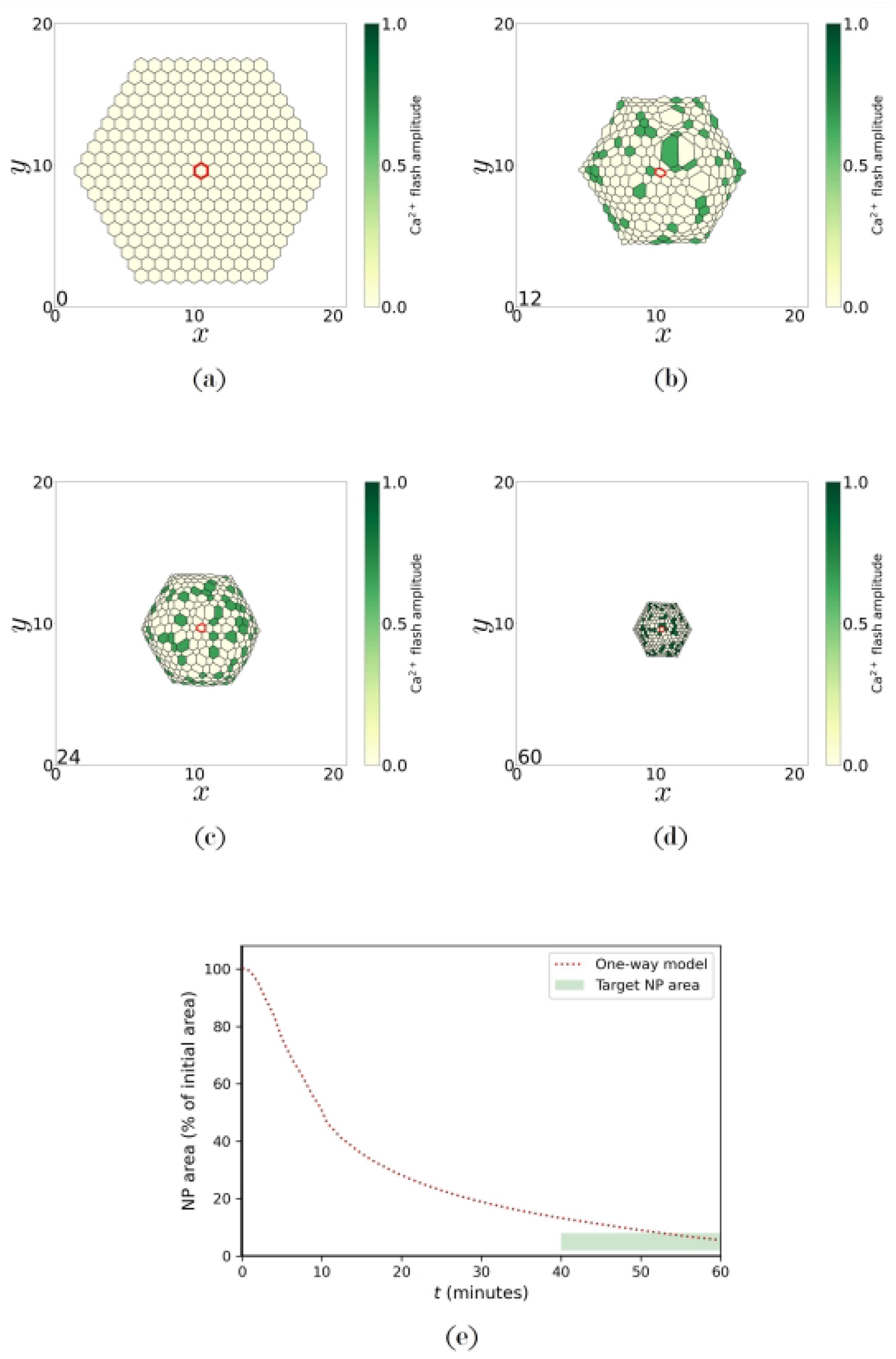
AC of the anterior NP as simulated by the one-way model. Snapshots of the apical surface of the anterior NP where yellow and green cells represent inactive and active cells, respectively, at: (a) *t* = 0, (b) *t* = 12, (c) *t* = 24, (d) *t* = 60 minutes, and (e) the evolution of the NP area over time. The shaded green region indicates the target range of 2%-8% for area contraction. The cell at the centre of the NP is marked with a red border. The impact of Ca^2+^ flashes on the area of the ‘marked’ cell can be seen in Fig 6.

**Table 1.** Parameter values used to generate Figs 4 - 8 and Fig S1.

| Parameter | Description | Value | Source |
| --- | --- | --- | --- |
| $\delta t$ | Timestep | 0.2 | [14] |
| $K_n^A$ (SE) | Area elasticity coefficient<br>(surface ectoderm cells) | 0.02 | [14, 43]* |
| $K_n^P$ (SE) | Perimeter elasticity coefficient<br>(surface ectoderm cells) | 0.01 | [14, 43]* |
| $T_{0n}$ (SE) | Baseline line tension<br>(surface ectoderm cells) | 0.01 | [14, 43]* |
| $K_n^A$ (ini) | Initial value of area elasticity coefficient<br>(NP cells) | 0.03 | [14]* |
| $K_n^P$ (ini) | Initial value of perimeter elasticity coefficient<br>(NP cells) | 0.02 | [14]* |
| $T_{0n}$ (ini) | Initial value of baseline line tension<br>(NP cells) | 0.01 | [14]* |
| $K_n^A$ (max) | Maximum value of area elasticity coefficient<br>(NP cells) | 0.3 | [14] |
| $K_n^P$ (max) | Maximum value of perimeter elasticity coefficient<br>(NP cells) | 0.2 | [14] |
| $T_{0n}$ (max) | Maximum value of baseline line tension<br>(NP cells) | 0.1 | [14] |
| $R_A$ | Growth rate of area elasticity coefficient<br>(NP cells) | 0.0003 | estimated |
| $R_P$ | Growth rate of perimeter elasticity coefficient<br>(NP cells) | 0.002 | estimated |
| $R_T$ | Growth rate of baseline line tension<br>(NP cells) | 0.002 | estimated |
| $A_n^0$ (ini) | Initial value of natural area<br>(all cells) | 0.785 | [14]* |
| $A_n^0$ (min) | Minimum value of natural area<br>(NP cells) | 0.015 | [9]* |
| $\kappa_A$ | Contraction rate of natural area<br>(NP cells) | 0.0008 | estimated |
| $\omega_A$ | Threshold parameter for the contraction of<br>natural area (NP cells) | 1 | estimated |
| $k_\xi$ | Scaling factor for the baseline tension<br>(NP cells) | 4 | estimated |
| $\tau$ | Activation period<br>(NP cells) | 67 | [11, 17]* |

\* These values were not directly taken from the cited studies; they were estimated using data presented in those studies.

**Fig 5.**
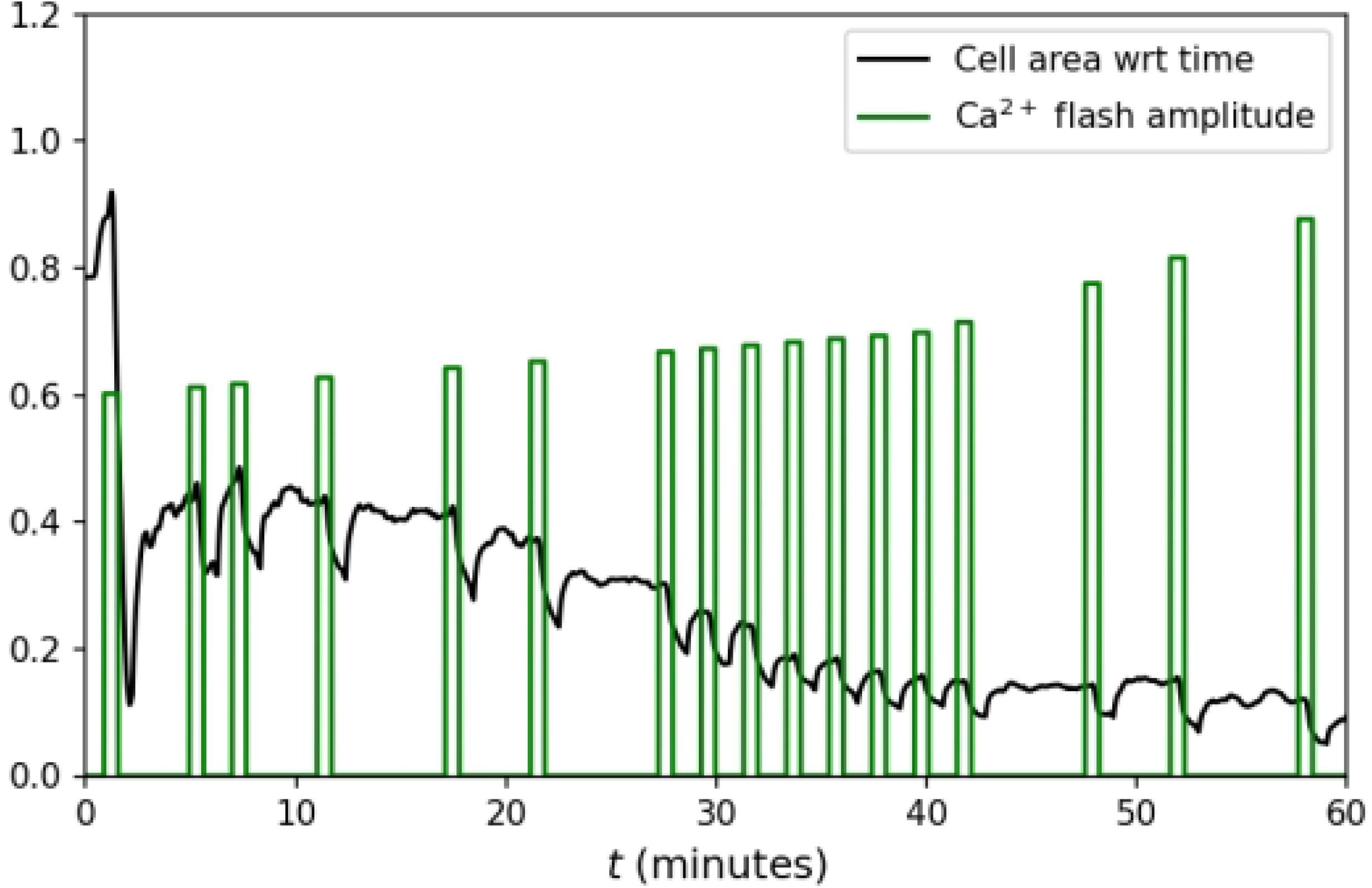
Final configuration of the simulated anterior NP and ectoderm (virtual cells). The apical surface of the anterior NP cells (solid yellow: inactivated, solid green: activated) and the virtual cells of the surface ectoderm (pink) in the one-way model, at the end of the 60-minute AC period.

**Fig 6.**
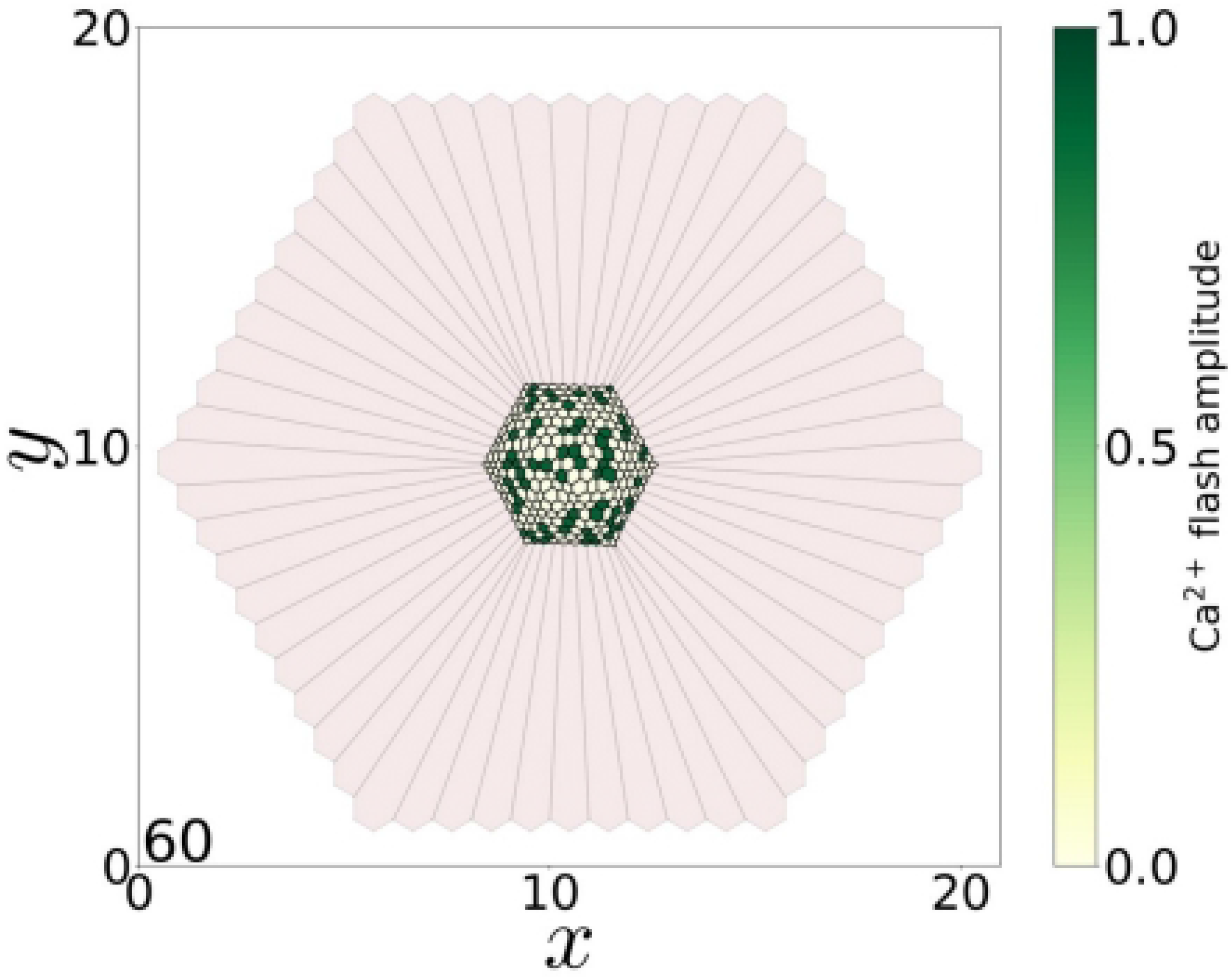
The effect of Ca^2+^ flashes on the apical surface area of a cell. Time evolution of the apical surface area of the ‘marked’ cell at the centre of the anterior NP visualised in Fig 4 with a red boundary.

**Fig 7.**
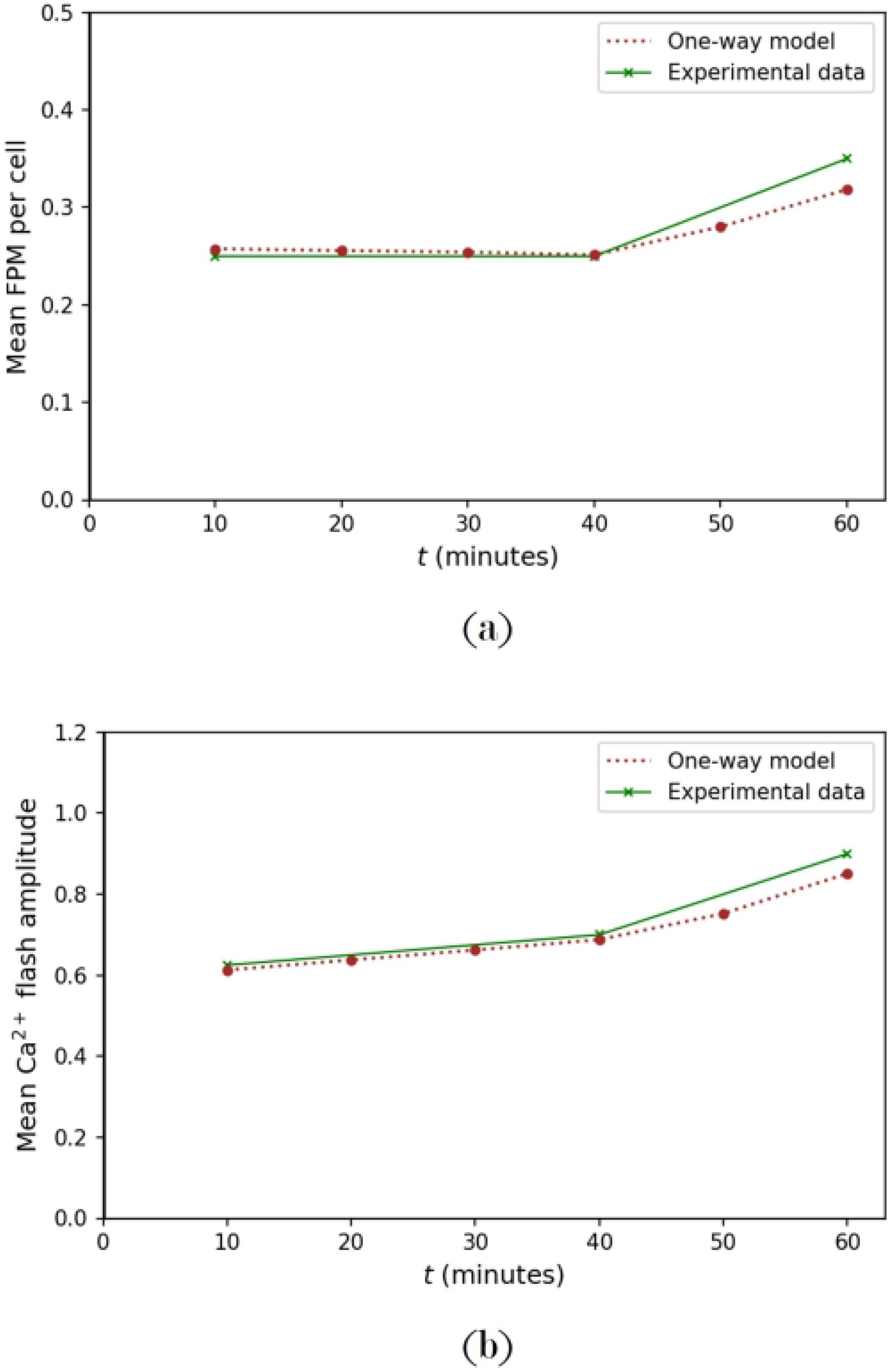
Evolution of the frequency and amplitude profiles of Ca^2+^ flashes in the one-way model: (a) mean number of flashes per minute (FPM) per cell, (b) mean amplitude of Ca^2+^ flashes. The experimental data (green) is input into the one-way model, which generates the corresponding frequency and amplitude responses (brown).

Fig 4(e) visualises the monotonic decrease of the surface area over time. We see that at *t* = 60 minutes, the area has constricted to 5.5% of its initial value, that is, the simulations agree with experimental findings [11, 18]. Fig 5 visualises the deformation of the virtual surface ectoderm cells at the end of the 60-minute AC period.

In Fig 6, we visualise the Ca^2+^ flashes and the apical surface area of the ‘marked’ cell (red boundary) in Fig 4, while the cell undergoes pulsed contractions. This reproduces experimental observations (see Fig 1c).

Figs 7a and 7b showcase the successful integration of Ca^2+^ frequency (Equation (9)) and amplitude (Equation (10)) profiles into the model. As AC progresses, both the frequency and amplitude of Ca^2+^ flashes increase as intended, consistent with experimental findings [11]. The measurements of FPM per cell and flash amplitude (averaged over all cells in the anterior NP) were taken over 10-minute time windows.

Fig 8 visualises the frequency distribution of the Ca^2+^ flashes at the 30 and 60 minute marks to demonstrate the spatial spread of the flashes mid-way and at the end of the AC process.

The one-way model also reproduces another key experimental observation: the continuous elevation of cytosolic Ca^2+^ inhibits the contraction of the apical surface of the anterior neural plate, leading to the failure of NTC [11]. When all anterior NP cells in the one-way model are continuously activated throughout the simulation, AC is effectively inhibited (see Fig S1). This highlights the critical importance of the Ca^2+^ refractory period and asynchronous Ca^2+^ flashes in the proper morphogenesis of the NP.

The one-way model is a stochastic model since the Ca^2+^ flashes are stochastic.

Therefore, to gain an accurate understanding of the model’s behaviour, we performed 100 realisations of the model, using the parameters in Table 1. We then calculated the range, mean, and standard deviation for the samples of the NP area, at every time step. The maximum standard deviation in NP area across the stochastic iterations was found to be less than 5%. Upon the conclusion of AC, all iterations converged to the same NP area. Given that there is very little variation in the NP area across the realisations, a single simulation of the model offers sufficiently accurate results.

Running the one-way model, it was found that both the adhesion force at the cell edges and the vertex damping function we have incorporated are essential for ensuring numerical stability; that is, not incorporating either of the two processes leads to numerical instability before the anterior NP contracts to the target area.

In summary, we have developed a one-way mechanochemical vertex model that replicates a wide range of experimental behaviours exhibited by the anterior NP during the AC phase of NTC. Table 2 compares the one-way model with the Suzuki model and highlights that the one-way model significantly improves the Suzuki model, reproducing more experimental findings.

**Fig 8.**
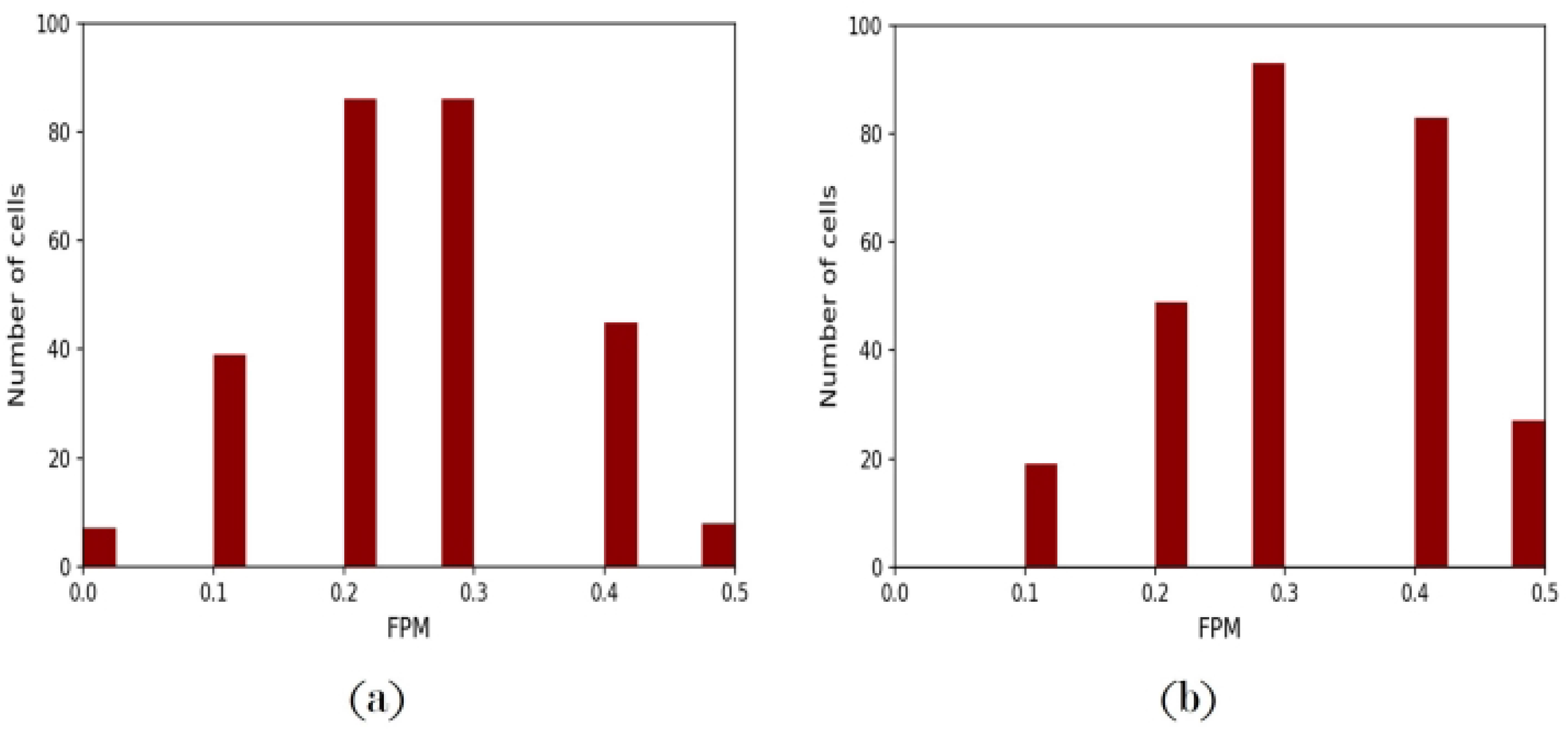
Frequency distribution of Ca^2+^ flashes over the anterior NP cells in the one-way model. Flashes per minute (FPM) per cell at: (a) 30 and (b) 60 minutes in Fig 7a.

**Table 2.** Comparison of the experimental findings reproduced by the Suzuki model and by the one-way model.

| Experimental findings | Suzuki model | One-way model |
| --- | --- | --- |
| Pulsed contractions at the cell level | Yes | Yes |
| Monotonic contraction at the tissue level | Yes | Yes |
| Incorporates effect of surface ectoderm | No | Yes |
| Anterior NP constricts to 2-8% of initial area | No | Yes |
| Latency between $\text{Ca}^{2+}$ flash and cell contraction | No | Yes |
| Frequency of $\text{Ca}^{2+}$ flashes increases with time | No | Yes |
| Amplitude of $\text{Ca}^{2+}$ flashes increases with time | No | Yes |
| NTC fails for sustained elevation of $\text{Ca}^{2+}$ | No | Yes |

### The two-way mechanochemical model

In the one-way model, the Ca^2+^ frequency and amplitude profiles were inputs of the model. To capture the two-way coupling between Ca^2+^ signalling and cellular mechanics [26–28, 42], we extend the one-way model to create a two-way mechanochemical model, where the Ca^2+^ frequency and amplitude profiles emerge as outputs.

We incorporate the feedback from cellular mechanics on Ca^2+^ signalling by introducing a ‘stretch activation’ mechanism into the model. Stretch-sensitive Ca^2+^ channels (SSCCs) on the cell membrane trigger the Ca^2+^-induced Ca^2+^ release (CICR) mechanism when the cytosol is stretched [22]. While there are continuum models that characterize the Ca^2+^ influx through SSCCs as being directly proportional to the dilatation of the cytosol [17, 22, 55], to the best of our knowledge, the two-way model presented here is the only vertex model that considers the effect of SSCC activation on cytosolic Ca^2+^ levels.

### Governing equations

The two-way model inherits Equations (1)-(4) and Equation (6) from the Suzuki model and Equations (12)-(23) from the one-way model. Equations (9) and (10) from the one-way model are not used since they correspond to the input Ca^2+^ frequency and amplitude profiles. Also, very importantly, in the one-way model, we replace Equation (11) for the mechanism for cell activation by Ca^2+^ with the SSCC-driven activation mechanism expressed by Equations (24)-(27)) below. Also, the two-way model revises modelling assumption (1.1), retains modelling assumptions (1.2)-(1.6), and introduces three new modelling assumptions (2.1)-(2.3) that justify Equations (24)-(27)).

### Probability of SSCC-driven cell activation

#### Assumption 1.1a.

*After the refractory interval in a Ca*^*2+*^ *flash elapses, the cell has an activation probability that depends on the number of stretch-activated edges in the cell and on the sensitivity of the SSCCs*.

#### Assumption 2.1.

*When an edge is stretched beyond a certain limit by the contraction of an adjacent cell, SSCCs on the edge open, leading to stretch-activation of the edge*.

#### Assumption 2.2.

*The sensitivity of the SSCCs increases with cortical tension which increases due to the accumulation of actomyosin in the cell’s apical cortex* [56–58]. *We assume that* 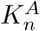 *is a measure of the level of cortical actomyosin*.

From studies on human retinal pigment epithelial cells [55] and on epithelial cells in the *Xenopus* NP [17], it is known that SSCCs on the cell membrane open in response to the dilatation or stretching of the cytosol; this may induce a Ca^2+^ flash within the cell. We hypothesize that ‘stretch activation’ serves to oppose the stretching of the cytosol by inducing a Ca^2+^ flash within the cell, to trigger cell contraction.

We define the probability of SSCC-driven cell activation, *p*_*n*_, as follows:

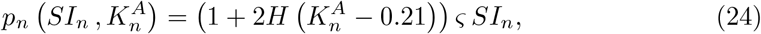

where *SI*_*n*_ is the stretch index of the cell, *ς* is the sensitivity of the SSCCs on a cell edge, 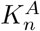 is the area elasticity coefficient, and *H* is the Heaviside function. The term 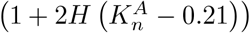 transitions from 1 to 3 at 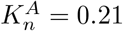 and represents a scaling function that scales the sensitivity of the SSCCs with 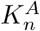.

Equation (24) is derived with reference to the frequency profile of Ca^2+^ flashes (Equation (9)), noting that they begin to increase at *t* = 4000, that is, at 40 minutes. At this time, an approximate value of 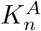 can be determined 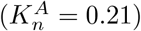. When this value is exceeded, the sensitivity of the SSCCs in a cell increases significantly. This behaviour is captured by the Heaviside function in Equation (24). The full derivation for the probability of SSCC-driven cell activation given by Equation (24) can be found in Appendix S1D.

Note that the change in SSCC sensitivity is likely best represented by a Hill function. However, for simplicity, we use the Heaviside function to model a step change from a low to a high value, at a specific actomyosin concentration - this avoids introducing an additional parameter for steepness.

The stretch index of the cell can range from 0 to 6, since we model the cells as hexagons, and is defined as follows:

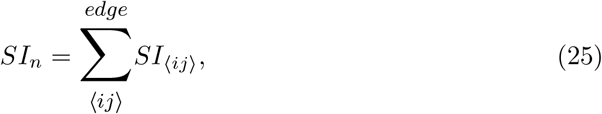

where *SI*_⟨*ij*⟩_, the stretch index of the cell edge, is defined as follows:

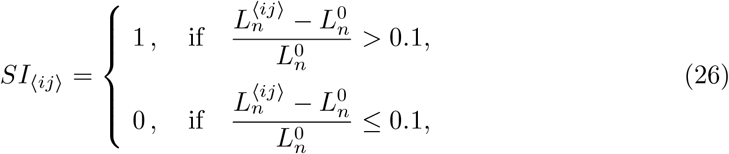

where 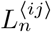 is the length of the cell edge and 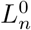 is the natural length of the cell.

We assume that the SSCCs are opened (edge activation) when the strain of a cell edge exceeds a predefined limit. In the absence of experimental data, we use 0.1 as an initial estimate for this strain limit. If an edge is activated in an inactive cell, the cell has a non-zero probability of experiencing a Ca^2+^ flash after every unit time interval. Equation (24) illustrates that the probability of cell activation increases with the number of activated edges in a cell. Conversely, if no edges are activated, the probability of cell activation is zero.

### Ca^2+^ flash amplitude

#### Assumption 2.3.

*The amplitude of the induced Ca*^*2+*^ *flash increases linearly with the level of cortical actomyosin in the cell*.

We define the amplitude of the induced Ca^2+^ flash, *A*, as follows:

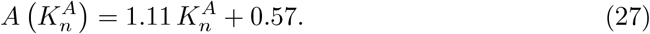

Since the relationship between 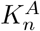 and Ca^2+^ flash amplitude is unknown, we assume a linear relationship, for simplicity. The Ca^2+^ amplitude profile (Equation (10)) indicates that *A*_(min)_ = 0.6 at the start of AC, when cortical actomyosin levels are at their lowest 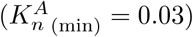, and *A*_(max)_ = 0.9 towards the end of AC, when cortical actomyosin levels are at their highest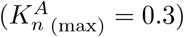. Equation (27) is a linear fit to this data.

To our knowledge, there are no studies examining the influence of cortical actomyosin enrichment on the opening of the SSCCs, the CICR mechanism and, consequently, on the amplitude of Ca^2+^ flashes. In the one-way model, we made the assumption that the amplitude of a Ca^2+^ flash determines the extent of myosin activity and, thereby, cell contraction. It is understood that the amplitude of Ca^2+^ flashes needs to increase to counteract the elastic forces arising from the accumulating cortical actomyosin and facilitate cell contraction. Therefore, we hypothesize that the accumulation of cortical actomyosin signals leads to increase in Ca^2+^ flash amplitude.

### Simulations

We implemented the two-way model in CelluLink by solving Equations (1)-(4), Equation (6), and Equations (12)-(27) for the parameter values in Table 3, using the same domain, boundary conditions, and initial tissue configuration as in the one-way model.

Unlike the one-way model, a fraction of cells needs to be activated at *t* = 0 to trigger the contraction of the NP. Therefore, an additional parameter, *α*, is introduced to represent the fraction of NP cells activated at *t* = 0. We estimate *α* = 0.1 based on insights from experimental observations of cell activation at the onset of AC during NTC [9, 11].

*In vivo*, this initial activation might stem from the mechanical forces generated during the convergent extension phase preceding AC [9, 11], or it could result from chemical signals such as Sonic Hedgehog, which plays a crucial role in regulating Ca^2+^ activity during neurulation

Fig 9 compares the monotonic contraction of the apical surface area of the anterior NP, as simulated by the one-way and by the two-way model. Both models exhibit a similar rate of constriction, down to 5.5% of the initial area.

In Figs 10a and 10b, we visualise the Ca^2+^ frequency and amplitude profiles which arise as outputs of the two-way model. The number of flashes per minute per cell and the average flash amplitude (averaged over all cells in the anterior NP) were taken over 10-minute intervals. Fig 11 visualises the distribution of FPM per cell in the two-way model, at 30 (left) and 60 minutes (right). We see that more cells exhibit higher FPM at 60 minutes, consistent with experimental observations.

Similarly to the one-way model, we performed 100 stochastic realisations of the two-way model, using the parameters listed in Table 3. The maximum standard deviation in the NP area across stochastic iterations was here also found to be less than 5%. Upon the conclusion of AC, all iterations converged to the same NP area. Given that there is very little variation in the NP area across stochastic realisations, data from a single iteration of the model offers sufficiently accurate results.

**Fig 9.**
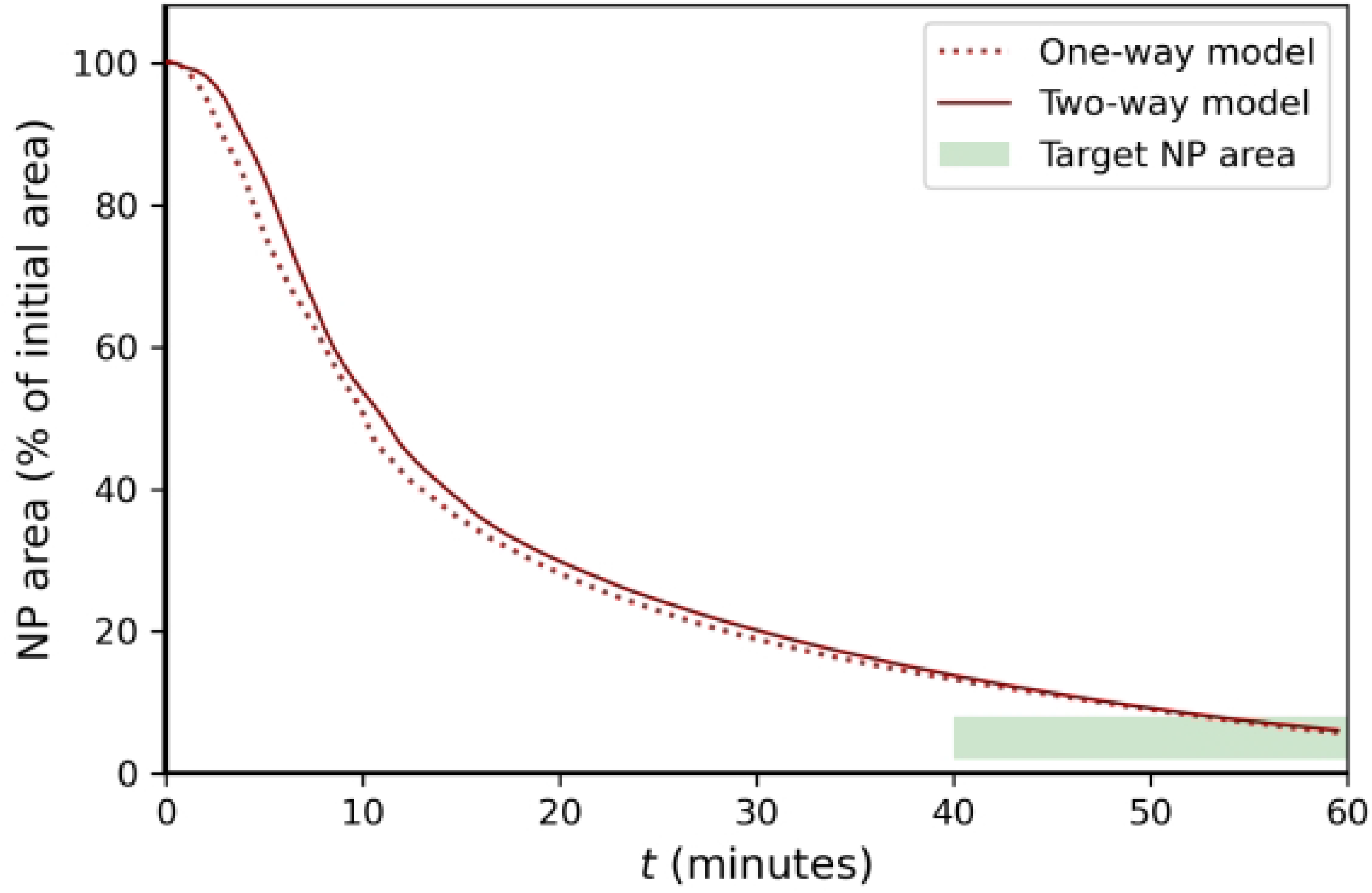
Time evolution of the apical surface area of the anterior NP, as simulated by the one-way and the two-way model. The reduction to 2-8% of the initial NP area observed in experiments is shown with the shaded green area.

**Table 3.**
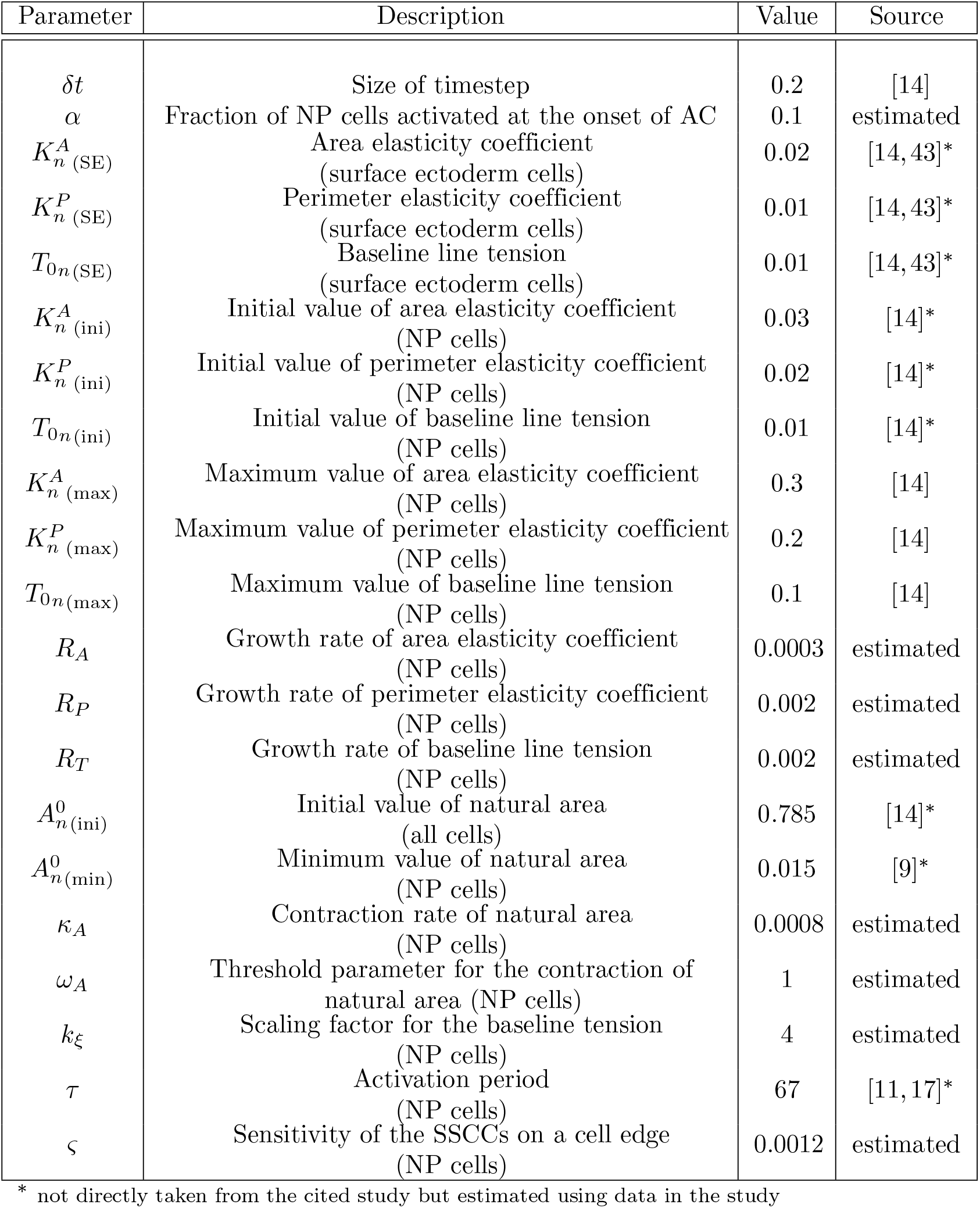
Parameter values used to generate Figs 9 - 11.

**Fig 10.**
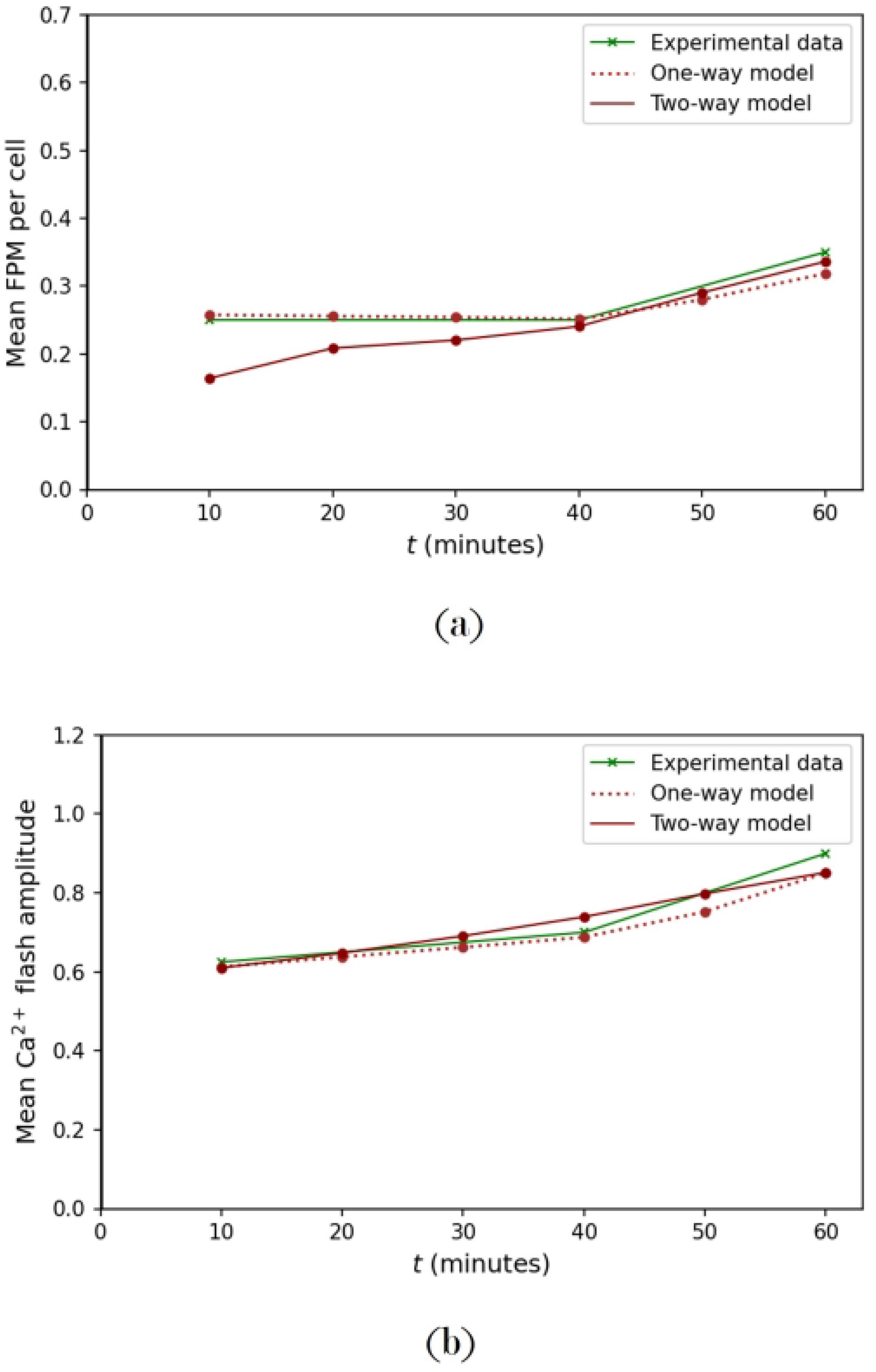
Frequency and amplitude profiles of Ca^2+^ flashes in the one-way and two-way models. Evolution of (a) mean number of flashes per minute (FPM) per cell, (b) mean amplitude of Ca^2+^ flashes, emerging as model outputs.

**Fig 11.**
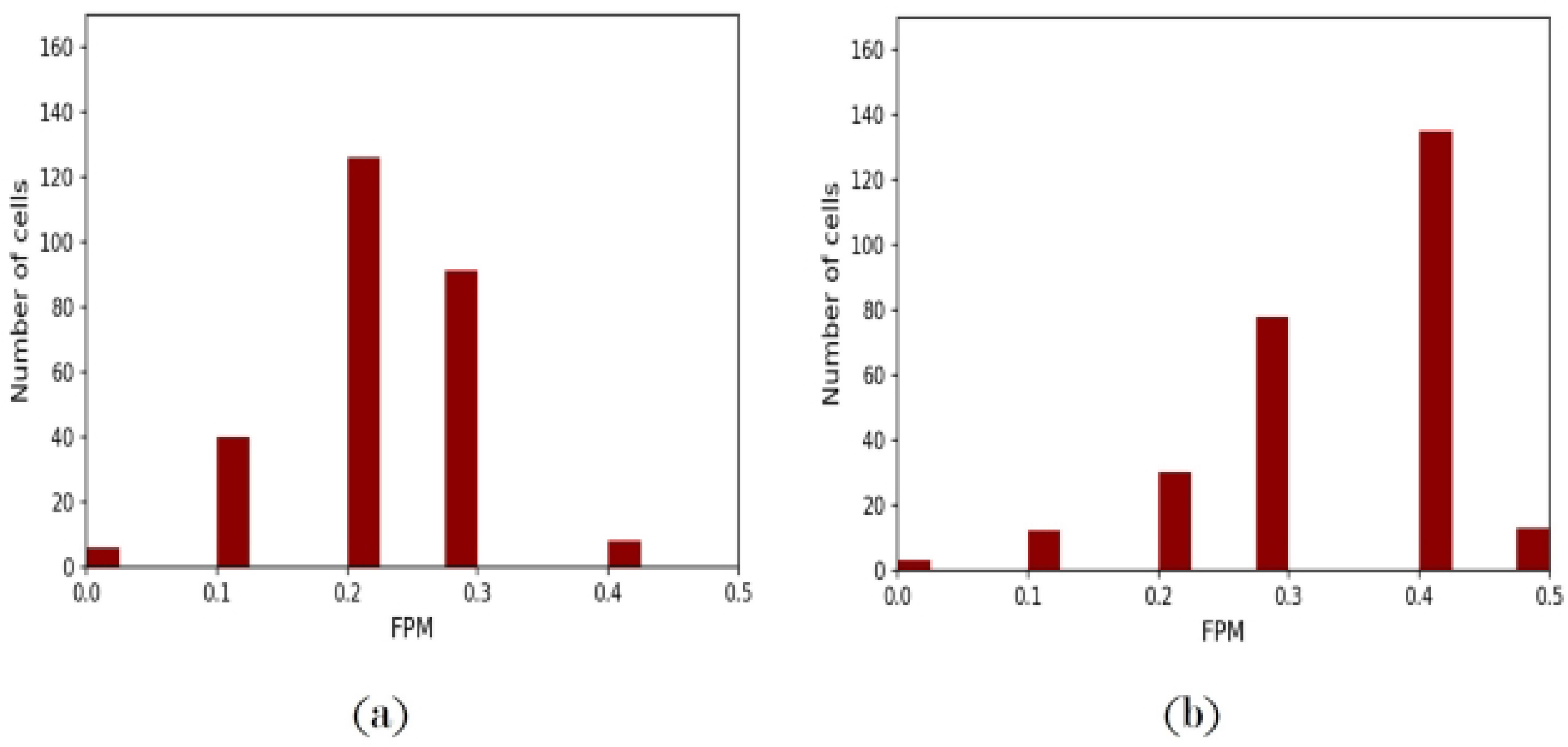
The frequency distribution of Ca^2+^ flashes over the anterior NP cells in the two-way model. Histograms depicting the distribution of flashes per minute (FPM) per cell at: (a) *t* = 30 and (b) *t* = 60 minutes in Fig 10a.

Table 4 compares the two-way model with the one-way model in terms of their ability to reproduce experimentally findings. The two-way model replicates all the results of the one-way model and reproduces additional behaviours that arise as a consequence of the stretch activation mechanism and the interaction between cells.

Note that the two-way mechanochemical coupling makes it possible to also capture short-range *multicellular Ca*^*2+*^ *transients* which have been observed in [11, 14]. In the case of a single-cell Ca^2+^ transient, a cell is spontaneously activated, as seen in the one-way model. In contrast, a multicellular Ca^2+^ transient involves the coordinated activation of multiple cells, where an activated cell triggers the activation of its neighbouring cells through mechanochemical transduction. This process allows a Ca^2+^ wave to propagate from cell to cell in the two-way model (see Video S1).

**Table 4.** Comparison of the experimental findings captured by the one-way and two-way models.

| Experimental findings | One-way model | Two-way model |
| --- | --- | --- |
| Pulsed contractions at the cell level | Yes | Yes |
| Monotonic contraction at the tissue level | Yes | Yes |
| Incorporates effect of surface ectoderm | Yes | Yes |
| Anterior NP constricts to 2-8% of initial area | Yes | Yes |
| Latency between $\text{Ca}^{2+}$ flash and cell contraction | Yes | Yes |
| Frequency of $\text{Ca}^{2+}$ flashes increases with time | Yes* | Yes |
| Amplitude of $\text{Ca}^{2+}$ flashes increases with time | Yes* | Yes |
| NTC fails for sustained elevation of $\text{Ca}^{2+}$ | Yes | Yes |
| Two-way coupling between $\text{Ca}^{2+}$ and mechanics | No | Yes |
| Multicellular $\text{Ca}^{2+}$ transients (across 2-5 cells) | No | Yes |

\* $\text{Ca}^{2+}$ frequency and amplitude profiles as inputs of the model

## Conclusions

We have developed two new mechanochemical vertex models for AC during NTC, significantly improving the Suzuki model [14]—the one-way model and the two-way model. The one-way model incorporates, for the first time, the effect of the surface ectoderm on the morphogenesis of the anterior NP during the AC phase of NTC. Furthermore, a damping function for the motion of the vertices, and an adhesion term for the energy function are incorporated, as motivated by experimental evidence. The damping coefficient increases with actomyosin concentration. Actomyosin concentration increases as actomyosin accumulates in the cell cortex [14, 18]. The adhesion force is assumed to be driven by the cell’s internal pressure and adherens junctions [49–51], which increases as cell edge length decreases. These new features are essential for capturing the underlying biology accurately and ensure cell and tissue integrity during AC and, correspondingly, numerical stability. Moreover, the mechanical parameters of the cells, 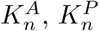, and 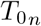, are assumed to grow with time.

In the one-way model, the Ca^2+^ flash frequency and amplitude profiles are inputs to the model. To capture the two-way coupling between Ca^2+^ signalling and cellular mechanics [26–28, 42], we have extended the one-way model to the two-way model by incorporating stretch sensitive Ca^2+^ channels (SSCCs) which enable the cell to sense mechanical stimuli and encode them into Ca^2+^ flashes. The two-way model reproduces all experimental findings exhibited by the one-way model but in the two-way model, the Ca^2+^ frequency and amplitude profiles emerge as outputs of the model. Moreover, the two-way model reproduces multicellular Ca^2+^ waves. We note that both models achieve a constriction down to 2-8% of the initial anterior NP area, in line with experiments. The full range of experimental behaviours captured by the models are summarised in Tables 2 and 4, respectively.

### Limitations and future work

Using the one-way model, the following experimental hypotheses are proposed, providing new directions for future research:

- Actomyosin exhibits persistent enrichment at cell-cell junctions, in contrast to the transient enrichment cycles followed by dissipation observed in medioapical actomyosin. This is essential for preserving cell and tissue integrity during AC.
- As cell-cell junctions decrease in size, apical adherens junctions exert a force, counteracting line tension, to prevent the cell surface from collapsing.
- The amplitude of a Ca^2+^ flash determines the extent of myosin activity and, thereby, cell contraction, during NTC.
- A refractory interval, characterized by a low Ca^2+^ level, is essential to alter the cell’s mechanical properties and activate the ratchet-like mechanism.

Similarly, using the two-way model, the following additional experimental hypotheses are proposed:

- The level of cortical actomyosin in a cell influences the amplitude of induced Ca^2+^ flashes, either by modulating the opening of the SSCCs or by impacting the Ca^2+^-induced Ca^2+^ release (CICR) mechanism.
- At low levels of cortical actomyosin, SSCC sensitivity remains fairly constant, but beyond a certain threshold of cortical actomyosin, SSCC sensitivity increases sharply.
- During AC, intercellular Ca^2+^ signalling between anterior NP cells occurs via mechanochemical transduction.
- A fraction of anterior NP cells must be activated to initiate AC. This initial activation could be caused by mechanical forces generated during convergent extension, or chemical signals.

In spite of the good agreement of the models with experiments, they have some limitations. Firstly, the NTC involves three-dimensional folding of the NP and, thus, 2D vertex models cannot fully capture NP morphogenesis. Specifically, the 2D models are unable to capture the spatial distribution of constriction foci on the anterior NP, regions of the anterior NP that display increased constriction under normal conditions during AC. Constriction foci are located along the periphery (hinge points) and in the medial region of the anterior NP. Sustaining high Ca^2+^ levels pharmacologically disrupts the normal constriction patterning, causing the distribution of foci to become random. This disruption leads to NTC failure and consequently to embryo defects. These results suggest that asynchronous and cell-autonomous Ca^2+^ flashes and contraction pulses are necessary for the correct temporal and spatial distribution of constriction and, as a result, for healthy neural tube closure [11].

To capture the spatial distribution of constriction, we considered modelling the constriction foci as regions of elevated cellular constriction. However, experimental data indicate that, as NTC progresses, all cells constrict to the same degree to promote tissue folding [9]. In our 2D vertex models, this would result in the same final tissue shape regardless of the initial spatial patterning of constriction, that is we cannot capture this morphogenetic behaviour with a 2D model. We postulate that either a 3D apical vertex model or a full 3D mechanochemical vertex model would capture the spatial distribution of constriction foci on the anterior NP.

Another limitation is that our models use simplified bistable Ca^2+^ dynamics, where a cell is either in a low Ca^2+^ (inactivated) state or a high Ca^2+^ (activated) state. More recent models of Ca^2+^ signalling incorporate more sophisticated Ca^2+^ dynamics, such as IP_3_-mediated Calcium-Induced Calcium Release (CICR) which has been validated by experiments [16, 17, 24]. Specifically, when a cell undergoes Ca^2+^ activation in our models, we assumed that the Ca^2+^ concentration remains elevated for a time, *τ*, before returning to its baseline value. In reality, the duration of cell activation is governed by CICR. Therefore, IP_3_-mediated Ca^2+^ dynamics could be incorporated as a next step.

This would be an important step in developing a model to assess the effects of potential treatments, such as clinical interventions. These interventions could involve pharmacologically increasing Ca^2+^ levels to induce cell contractions in regions of the NP that do not contract sufficiently or suppressing elevated Ca^2+^ levels in regions contracting too rapidly.

During NTC, two types of multicellular Ca^2+^ transients have been observed in the anterior NP: short-range Ca^2+^ waves that propagate over two to five cells, and long-range Ca^2+^ waves that propagate over tens of cells [11]. The two-way model successfully reproduces short-range Ca^2+^ waves but not long-range Ca^2+^ waves. It may be worth investigating whether more sophisticated Ca^2+^ dynamics, such as IP_3_-mediated CICR, could generate long-range waves.

Experimental validation of parameter choices remains a significant challenge for vertex models. Currently, there is no methodology to relate measured tissue properties (continuum-level parameters), such as Young’s modulus and traction stress, to the mechanical parameters of the vertex model (cell-level parameters). Prior studies have analytically derived expressions for the shear and bulk moduli of the tissue in their vertex models [60–62] and parametrised simulations to match experimental results [62]. The two-way model could be parametrised using a similar approach. Alternatively, conducting *in silico* stress tests could elucidate the relationship between cell-level and continuum-level parameters. For example, Honda and Nagai [63] explored the viscoelastic properties of a cell aggregate through a stress test on their vertex model. By analysing the deformation curves obtained from the stress test, they were able to determine the viscosity and elastic constants of the cell aggregate. A similar method could be employed to establish a connection between the properties of the anterior NP and the cell-level parameters in the two-way model.

To conclude, we have made a significant contribution by developing the first 2D mechanochemical vertex model that captures the two-way coupling between Ca^2+^ signalling and cellular mechanics, laying the foundation for future work.

## Supporting information

**Appendix S1**. Supplementary sections on parameter estimation (S1A) and derivations of the adhesion term (S1B), damping function (S1C), and probability of SSCC-driven cell activation (S1D).

**Fig S1. The effect of the continuous elevation of cytosolic Ca**^**2+**^ **on AC**. Snapshots of the apical surface of the anterior NP (green cells represent active cells) in the one-way model at: (a) *t* = 0, (b) *t* = 60 minutes, and (c) the evolution of NP area. The shaded green region indicates the target range of 2%-8% for area contraction. The continuous elevation of cytosolic Ca^2+^ inhibits the contraction of the apical surface of the anterior neural plate, resulting in the failure of NTC.

**Video S1. Time evolution of the apical surface of the anterior NP in the two-way model**. In the two-way model, the contraction of a cell can induce a Ca^2+^ transient in its neighbours by the ‘stretch activation’ mechanism. This process facilitates the propagation of a Ca^2+^ wave over a small group of cells.

